# A sample-to-report solution for taxonomic identification of cultured bacteria in the clinical setting based on nanopore sequencing

**DOI:** 10.1101/752774

**Authors:** Stefan Moritz Neuenschwander, Miguel Angel Terrazos Miani, Heiko Amlang, Carmen Perroulaz, Pascal Bittel, Carlo Casanova, Sara Droz, Jean-Pierre Flandrois, Stephen L. Leib, Franziska Suter-Riniker, Alban Ramette

## Abstract

Amplicon sequencing of 16S rRNA gene is commonly used for the identification of bacterial isolates in diagnostic laboratories, and mostly relies on the Sanger sequencing method. The latter, however, suffers from a number of limitations with the most significant being the inability to resolve mixed amplicons when closely related species are co-amplified from a mixed culture. This often leads to either increased turnover time or absence of usable sequence data. Short-read NGS technologies could address the mixed amplicon issue, but would lack both cost efficiency at low throughput and fast turnaround times. Nanopore sequencing developed by Oxford Nanopore Technologies (ONT) could solve those issues by enabling flexible number of samples per run and adjustable sequencing time. Here we report on the development of a standardized laboratory workflow combined with a fully automated analysis pipeline *LORCAN* (Long Read Consensus ANalysis), which together provide a sample-to-report solution for amplicon sequencing and taxonomic identification of the resulting consensus sequences. Validation of the approach was conducted on a panel of reference strains and on clinical samples consisting of single or mixed rRNA amplicons associated with various bacterial genera by direct comparison to the corresponding Sanger sequences. Additionally, artificial read mixtures of closely related species were used to assess *LORCAN*’s behaviour when dealing with samples with known cross-contamination level. We demonstrate that by combining ONT amplicon sequencing results with *LORCAN*, the accuracy of Sanger sequencing can be closely matched (>99.6% sequence identity) and that mixed samples can be resolved at the single base resolution level. The presented approach has the potential to significantly improve the flexibility, reliability and availability of amplicon sequencing in diagnostic settings.

## Introduction

The introduction of MALDI-TOF (matrix-assisted laser desorption/ionization time-of-flight) mass spectrometry has revolutionized routine microbiological diagnosis as it provides fast and reliable identification of clinically-relevant microorganisms [1]. The technology is particularly useful for species identification of bacterial isolates from diverse taxonomic groups [2,3], for which conventional methods or time-consuming biochemical tests need to be applied [2,4]. Although a large number of diagnostic laboratories in both industrialized and non-industrialized countries have adopted the new technology over conventional biochemical tests or molecular techniques for their routine applications, the method may be limited by several factors: First, the approach relies on the chemical composition of the microbial cell wall, and may thus be directly influenced by sample impurities or by the extraction method. Extraction is also difficult to standardise and often lead to difficulties to obtain valuable prepared samples resulting in suboptimal or lack of identification [5,6]. Second, sub-optimal identification may be caused by database limitations, including insufficient representation of reference species profiles in available commercial databases, absence of newly discovered species, and the existence of several commercial systems [6-8]. Third, MALDI-TOF is intrinsically a phenotyping/chemotyping method and is therefore indirectly related to the phylogeny of the organisms, even though most of the key proteins it uses consist of ribosomal proteins which are known to be highly valuable in phylogenetic studies [9,10].

Although the sequencing of the 16S rRNA gene is essential in the studies that describe the diversity of the human microbiome [11,12], the frequency of the use of 16S sequencing for species identification in clinical laboratories is decreasing [13], despite the usefulness of 16S rRNA gene sequencing to provide taxonomic classification for isolates that do not match recognized biochemical profiles, that only produce low identification score according to commercial systems, or that are not typically associated with human pathogens [13,14]. In the clinical microbiology laboratory, amplicon sequencing of 16S rRNA gene mostly relies on the Sanger sequencing method, which is based on chain termination via fluorescently labelled deoxyribonucleotides (dNTPs), capillary electrophoresis and fluorescence measurement [15]. Although the Sanger method is still the gold-standard for validating the accuracy of sequences from specific genes, when compared to more recent technologies, the method has a number of significant shortcomings: During a sequencing run, each capillary is limited to the production of one single sequence with a maximal length of about 1000 bp [16], resulting in low throughput, and high sequencing costs. Furthermore, the sequencing machines are comparably large and require maintenance, limiting their suitability for all types of laboratory settings. The most important limitation of the Sanger method is, however, its inability to resolve mixed amplicons [17]. Under routine diagnostic conditions, this frequently leads to either increased turnover time or lack of results [18], leading to potential delays or inaccuracies in patient treatment and management.

Recent sequencing technologies (i.e. second-generation sequencing technologies, such as provided by Illumina) might overcome most of these limitations, but are difficult to scale down in terms of turnover time and sequencing costs, especially when only few samples need to be sequenced at infrequent periods, making them inappropriate for the identification of bacterial isolates based on their 16S amplicon sequences in most diagnostic facilities. The third-generation single-molecule sequencing technology provided by Oxford Nanopore Technologies (ONT) may offer the necessary flexibility in throughput and is capable of producing reads with lengths of several hundred to several hundred-thousand bases at competitive costs [19]. Furthermore, ONT sequencers are small devices, virtually maintenance free and affordable for independent laboratories. Despite the constant improvement over the last years in read accuracy (with current read accuracy of about 96%), the remaining sequencing errors in single nanopore reads do not yet allow for an analysis at the read level. *De novo* assembly or consensus generation from the individual reads are therefore commonly used to generate sequences that are virtually free from substitution errors [20]. Additionally, polishing tools can be applied to remove remaining non-random errors such as indels in homopolymer regions [20-23]. Resulting sequences can then be directly substituted to Sanger sequences in existing classification pipelines or, due to the added flexibility in read length, may provide far higher resolution if the analyses are based on full-length marker genes or entire operons [24]. One obstacle for a broad adoption of nanopore sequencing in routine diagnostic laboratories is the added bioinformatic complexity as compared to established Sanger sequencing workflows. Furthermore, available workflows are often limited to the analysis of pure amplicons [20-23], include complex modifications of the ONT laboratory workflows [25,26], or lack published validation by using samples other than mock communities [27,28].

Here, we developed a complete workflow based on standard ONT protocols (supplemented with additional controls) and a fully automated analysis pipeline *LORCAN* capable of producing high-quality consensus sequences and thorough taxonomic analysis from pure and contaminated samples. Analysis parameters were evaluated with amplicons generated from reference strains of well-known pathogenic genera (*Bacteroides, Eggerthella, Enterococcus, Klebsiella, Mycobacterium, Campylobacter, Pseudomonas*) and validated on bacterial cultures obtained from patient material over several months. Furthermore, we explored the robustness of *LORCAN’s* consensus generation and species identification by analysing artificial mixtures of reads at different levels of genetic distances.

## Methods

### Samples, DNA extraction, PCR amplification

Bacterial isolates all originated from the Institute for Infectious Diseases (IFIK; Bern) Biobank. The IFIK provides the entire spectrum of medical microbiological diagnostic services to the largest Swiss hospital group (Inselgruppe) and other regional hospitals. The diagnostic division of IFIK (clinical microbiology) is ISO/IEC 17025 accredited to perform routine bacterial diagnostics from clinical samples. ATCC strains were obtained from LGC Standards (Wesel, Germany) and were grown on solid media as recommended by the manufacturer.

Overnight-grown bacterial cultures were harvested from agar plates and dissolved in 300 μl of Tris-EDTA (pH 8.0). DNA was extracted with a NucliSense Easymag (bioMérieux, Switzerland) robot according to the manufacturer’s protocol. 16S rRNA gene PCR was performed with the primer sets 16S_ f: 5′-AGAGTTTGATCMTGGCTCAG-3′ and 16S_ r: 5′-TACCGCGGCWGCTGGCACRDA-3′ (general bacteria) and mbak_f: 5′-GAGTTTGATCCTGGCTCAGGA-3′ and mbak_r: 5′-TGCACACAGGCCACAAGGGA-3′ (*Mycobacteria)* supplemented with the universal tails 5′-TTTCTGTTGGTGCTGATATTGC-3′ (ONT forward primer), 5′-ACTTGCCTGTCGCTCTATCTTC-3′ (ONT reverse primer), 5′-TGTAAAACGACGGCC AG-3′ (M13f, Sanger forward primer) or 5′-CAGGAAACAGCTATGAC-3′ (M13r, Sanger reverse primer). PCR reactions (25 μl) for general bacteria and *Mycobacteria* were assembled with 1 / 2.5 ng template, 10 μl of a 1.25 / 2.5 μM primer working solution and 12.5 μl Q5 Master-Mix. Amplification was performed in a GeneAmp 9700 Thermocycler (Thermo Fisher Scientific Inc., MA, USA) with the following program: 98°C for 1 min; 30 cycles of: 98°C for 10 s, 63°C for 15 s, 72°C for 30 sec; 72°C for 2 min. PCR products were purified with CleanNGS beads (CleanNA, Waddinxveen, NL) according to the manufacturer’s instructions with the following modifications: After the washing step an additional 3 sec centrifugation step was introduced and the purified DNA was eluted in 80 μl of Tris-HCI (0.01M, pH 8.0). Fragment size of the amplicons was analysed using the TapeStation D1000 assay (Agilent, Santa Clara, CA USA), concentrations were measured with the Qubit dsDNA BR assay (Thermo Fisher Scientific), and the purity of the DNA was analysed with a Nanodrop spectrophotometer (Thermo Fisher Scientific). Samples with DNA concentrations <1.05 nM were excluded from the analysis.

### Library preparation

A typical library consisted of the pooling of 2 to 15 clinical samples and 1 positive control (*Mycobacteria intracellulare*, amplified with general bacterial primers). Library preparation was performed with the kits EXP-PBC096, SQL-LSK109 (Oxford Nanopore Technologies, OX, UK), and using the supplementary reagents NEBNext End repair/dA-tailing Module (E7546, New England Biolabs, ON, CA), NEB Blunt/TA Ligase Master Mix (M0367, New England Biolabs), Taq 2X Master Mix (NEB M0270, New England Biolabs), CleanNGS beads (CleanNA). All modifications made to the manufacturer’s protocol (PCR barcoding (96) genomic DNA, PBAC96_9069_v109_revK_14Aug2019) are described in the following section (for a detailed protocol see Supplementary Text S1): AMPure beads were substituted with CleanNGS beads and the Hula-Mixer (Thermo Fisher Scientific) parameters “Orbital: 40 rpm, 07 s; Reciprocal: 89 deg, 2 s; Vibro: 5 deg, 2 s; Vertical position” were used. Barcoding-PCR reactions (12 cycles) were set up with 25.2 nmol of template per reaction. Raw barcoded PCR products were quantified with the Qubit dsDNA BR assay and pooled at equal molar proportions. If the total amount of DNA in a pooled library was below 9.23 pmol, “place-holder” (filling) barcoded samples were added to the pooled library to avoid flow cell underloading (see example of calculations and adjustments in Supplementary Text S1). Place-holder libraries were produced in advance from the same template as the positive controls, with 15 instead of 12 barcoding PCR cycles. Resulting PCR products were quantified with Qubit and stored at -20°C. The pooled library was purified (CleanNGS beads, 50 μl elution volume), and quantified with the Qubit dsDNA BR assay. The purified library pools were diluted to 140 nM before proceeding to the “End Preparation” step of the protocol.

### Sequencing

ONT-sequencing was performed on a GridIONX5 instrument (Oxford Nanopore Technologies) with real-time base calling enabled (*ont-guppy-for-gridion* v.1.4.3-1 and v.3.0.3-1, fast base calling mode). Sequencing runs were terminated after production of 1 million reads or when sequencing rates dropped below 20 reads per second. Purified PCR products were submitted to Sanger sequencing at Microsynth (Balgach, Switzerland).

### Bioinformatic analyses

#### LORCAN pipeline description

*LORCAN* was developed to facilitate reproducible ONT sequencing based marker gene analysis in diagnostics facilities. The pipeline written in *Perl* 5, R and *BASH* and runs on Linux servers. The code is publicly available [29] and is based on publicly available, third-party dependencies (Table S3). Major steps of the workflow are described in the following section (numbers correspond to the steps in Figure 2): Step 1) Basecalled reads are demultiplexed and adapters trimmed (*Porechop* [30], parameters: --format fasta, --discard_unassigned, --require_two_barcodes). Step 2) Reads are filtered by length, keeping only those with lengths of -20 to +100 bases around the modal sequence length (custom *Perl* and *R* scripts; Figure 2). Step 3) Reads are mapped to a non-redundant reference database (*minimap2* [31]; see database preparation below). Step 4) Reads are extracted, binned by taxonomic level (here species) and remapped to the reference sequence that obtained the highest number of mapped reads among all sequences of the corresponding species (*minimap2, SAMtools* [32], *SeqKit* [33]). Step 5) Consensus sequences are derived using a 50% majority rule consensus. Step 6) The 10 closest reference sequences are selected by sequence similarity to the consensus sequence (BLASTN, *BLAST+*, [34]). Step 7) Phylogenetic trees for each consensus sequence with its 10 closest references are created (*MAFFT* [35] with parameters *-maxiterate 1000 – localpair*; *Gblocks* [36] with parameters *-t=d, IQ-TREE* [37] with parameters *-m GTR+I+G -bb 1000 -czb*). Parameters of all software are also provided in the *LORCAN* GitHub repository.

**Figure 1.**
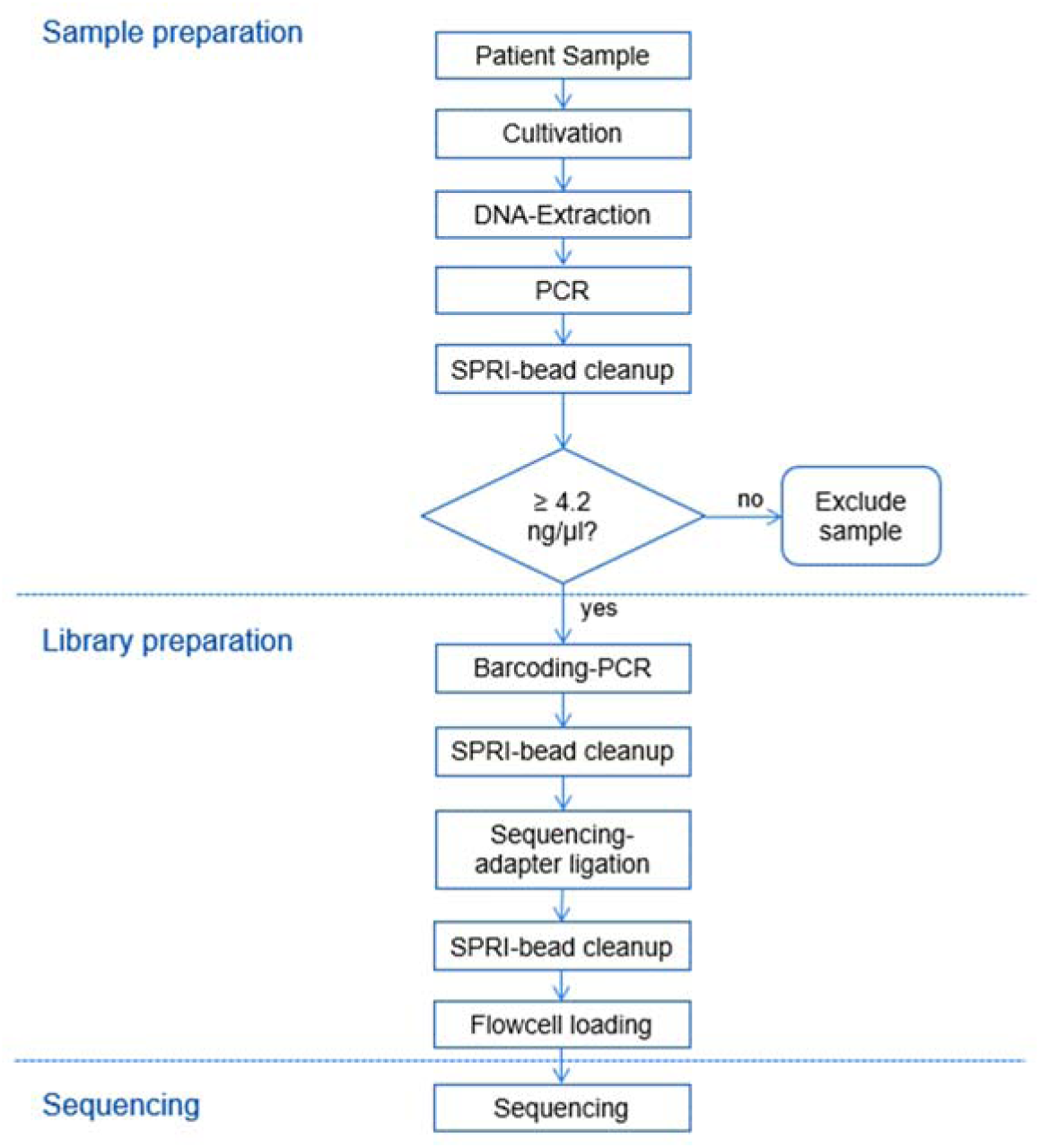
Overview of the wet laboratory workflow.

**Figure 2.**
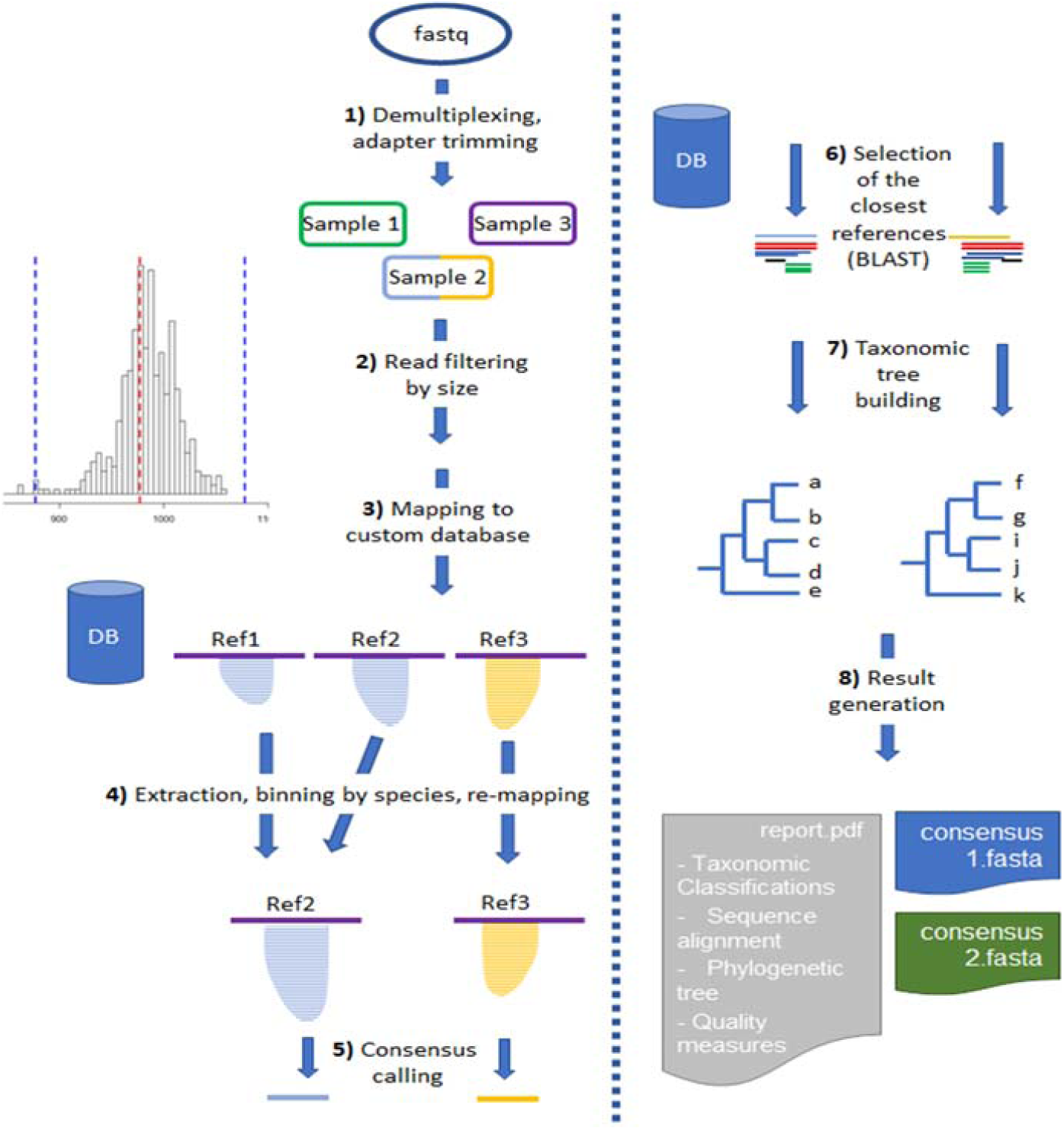
Steps involved in the *LORCAN* analysis pipeline. Step 1: “Sample 2” represents a mixed sample composed of two species; Step 3: Ref1 and Ref2 represent reference sequences from two different strains of the same species.

#### Database preparation

Reference databases used by *LORCAN* are non-redundant and assay specific. Detailed instructions for database creation are provided online at: https://github.com/aramette/LORCAN/. In short, the reference database (in this study: leBIBI SSU-rDNA-mk37_stringent, https://umr5558-bibiserv.univ-lyon1.fr/BIBIDOCNEW/db-BIBI.html; [38]) was trimmed to the region of interest (amplified region minus primers) and de-replicated (*Mothur* [39]), and sequence names were simplified (custom *Perl* scripts). The names of identical sequences are saved to a file during the dereplication step. The resulting non-redundant database is then used to generate a custom *BLAST* database which is used in *LORCAN* pipeline.

#### Sanger sequence analyses

Forward and reverse sequences were assembled into consensus sequences using *SeqMan Pro* (DNAStar, Madison, WI, USA), primers were trimmed manually, and ambiguous bases were resolved based on visual inspection of the chromatograms. Consensus sequences were taxonomically classified using the online tool *leBIBI QBPP* [38,40].

### Effects of parameter modifications on *LORCAN* results

#### Read length vs. consensus quality

Read sets were collected from a *LORCAN* output directory (output file 1_fasta/BC* .fasta produced by step 1; Figure 1) subsetted by size (50 reads per window, overlapping windows, width: 10 bases, offset: 2 bases) and fed back into *LORCAN* (steps 3 to 8; Figure 1). Consensus quality parameters (i.e. consensus length, fraction of reads of a sample used for generation of the top consensus sequence, numbers of gaps and numbers of ambiguous bases) were extracted from the *LORCAN* reports and visualized (*R* v.3.6.0, *RStudio* v.1.2.1335, *ggplot2*).

#### Numbers of read vs. consensus quality

Size selected reads sets produced from seven ATCC strains were collected from the *LORCAN* output directory (output file 1_fasta/BC* .mode_closest.fasta, produced by step 2; Figure 1) and subsampled to produce 100 read sets composed of 10 to 1000 reads each. *LORCAN* was run and quality parameters were extracted from the resulting *LORCAN* reports. Additionally, each top consensus sequence was compared to the consensus sequence produced from the full dataset (*BLAST*+ v. 2.6.0 default parameters).

#### SNV discrimination and performance with mixed samples

Size selected reads produced from 4 pure *Mycobacterium* amplicon samples were collected from the *LORCAN* output directories (output file 1_fasta/BC* .mode_closest.fasta, produced by step 2; Figure 1) and used to assemble artificial read sets, each composed of different proportions of two of the original samples (51 read set per sample pair, proportions of each species 0-100%, *Seqtk subseq* v.1.3-r106, https://github.com/lh3/seqtk). Reads were fed back into *LORCAN* and detected species compositions were extracted from the resulting *LORCAN* reports. Sequence identities of the paired species were determined based on pairwise alignment of the region flanked by the used primer sets (*Multalin*, version 5.4.1, http://multalin.toulouse.inra.fr/multalin/; [41]).

#### Influence of database completeness on consensus accuracy

A set of 7 ATCC reference strains was sequenced and analysed with *LORCAN* using the full non-redundant leBIBI 16S rRNA database generated for the general bacterial primer set). The resulting top consensus sequences were extracted, combined with the above-mentioned database, the resulting dataset was aligned (*MAFFT* v7.313, FFT-NS-1, progressive method) and pairwise distances were calculated (*Mothur* v. 1.40.5, *dist.seqs*, calc=eachgap, countends=F, cutoff=0.20). For each consensus sequence 10 subsets of sequences with minimal distances below thresholds between 0 and 0.1 were extracted (*Seqtk subseq*), and minimal distances between each dataset and the corresponding consensus sequence were analysed (x-axis in Figure 6). The seven read sets (ATCC strains) were re-analysed with *LORCAN* and the corresponding subsetted databases and the resulting consensus sequences were extracted. Top consensus sequences from each sample-database combination were extracted, combined with the consensus sequences generated with the full database, and aligned (*MAFFT* v7.313, L-INS-I, iterative refinement method (<16) with local pairwise alignment information). Pairwise distances were analysed as described above and distances between the consensus sequences generated from the full and the subsetted databases were extracted (y-axis in Figure 6).

**Figure 3.**
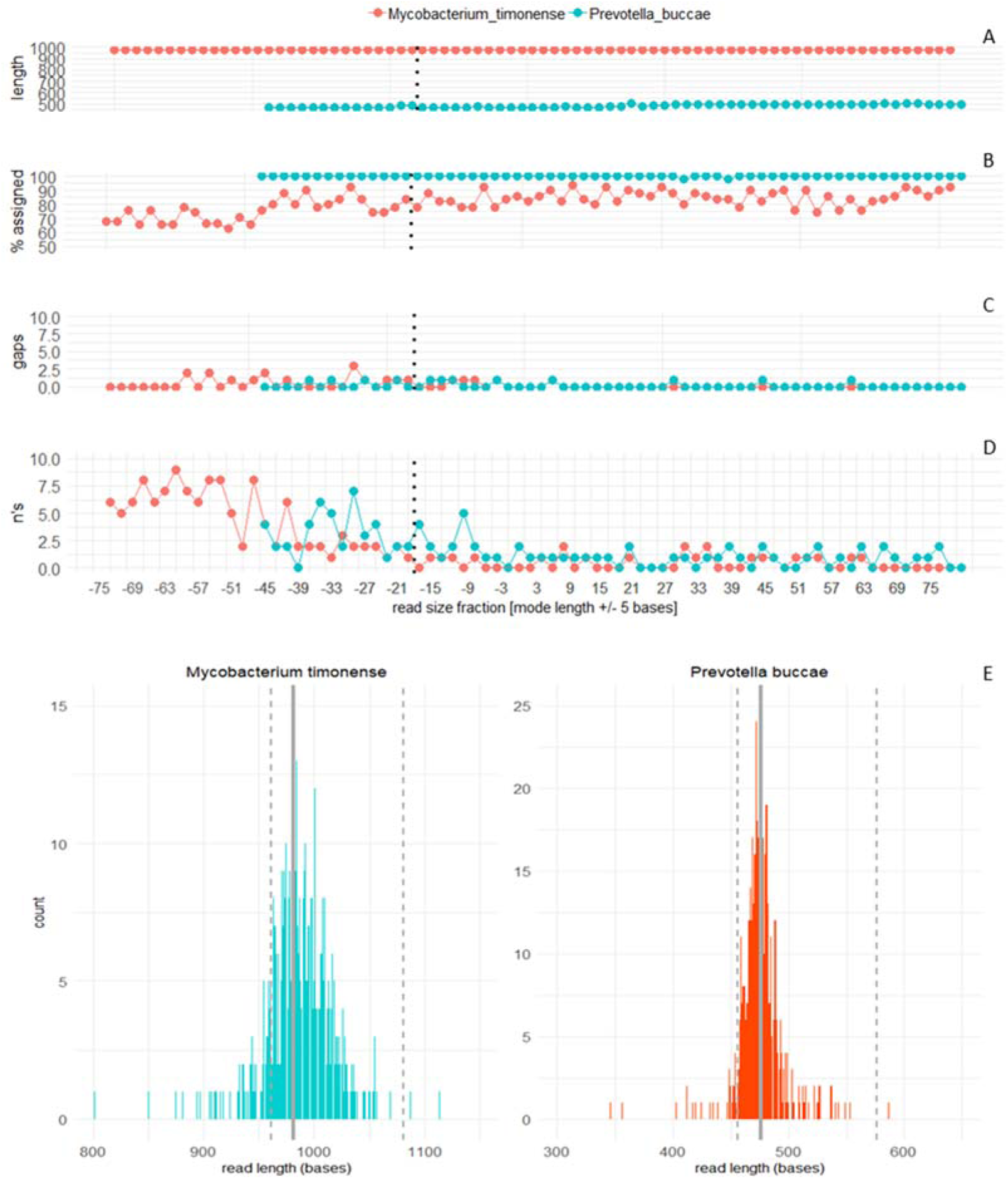
Influence of read size fraction on the quality of the *LORCAN* consensus sequences. **A**) consensus length. **B**) fraction of the analysed reads used for consensus generation. **C**) and **D**) numbers of gaps and ambiguous sequences in the consensus sequences. The x-axis represents differences to the modal size of the complete read set and the centre of the 11-base size windows used for read subsetting. Dotted lines indicate the recommended lower cut-offs for size selection. Missing points are a result of insufficient numbers of reads (<50 reads) in the respective size fraction. **E**) Size distribution of the raw FASTQ reads and recommended size thresholds. Solid lines represent the modes of the read length distributions. Dotted lines indicate the recommended lower and upper cut-offs for size selection.

**Figure 4.**
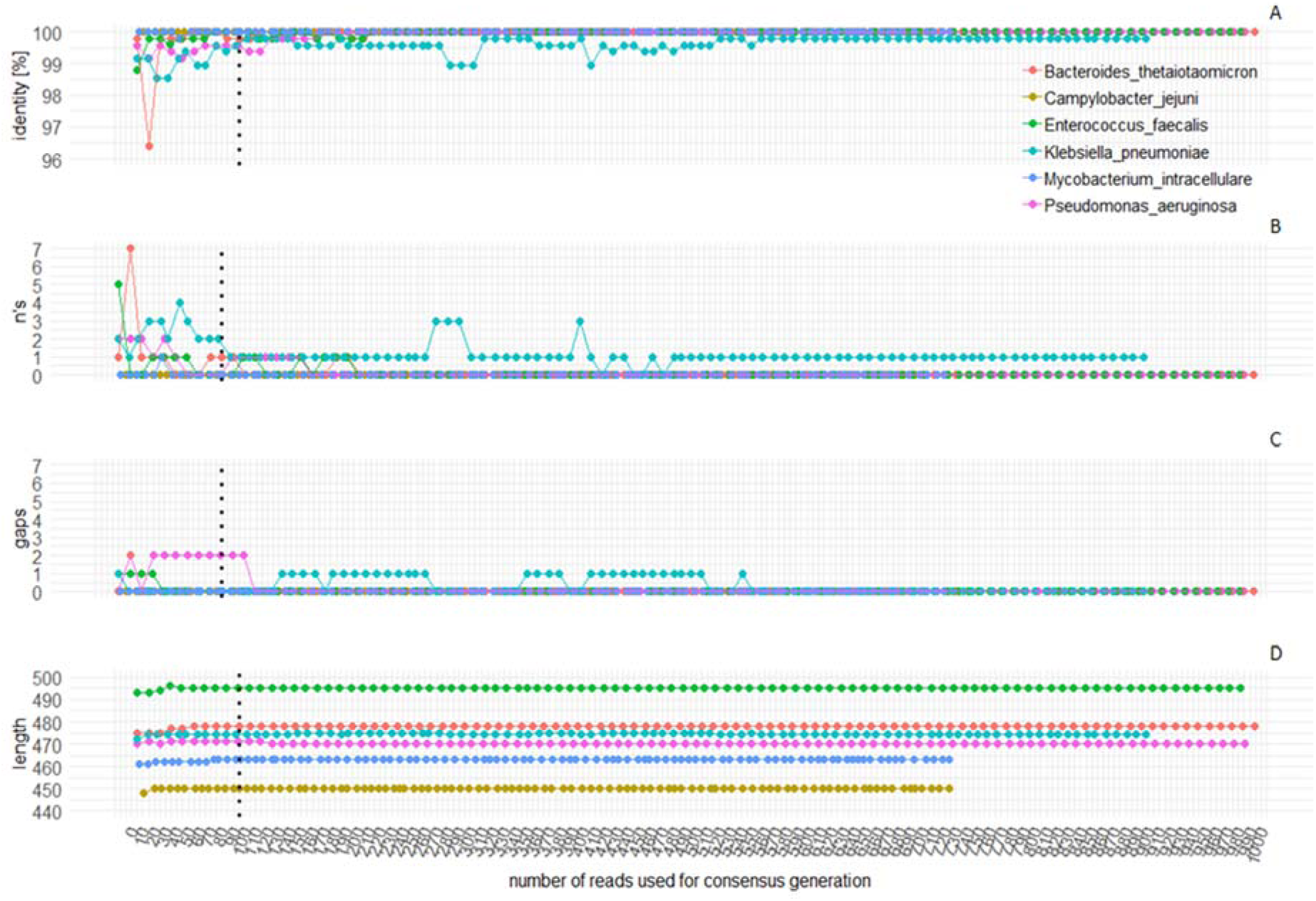
Influence of input read number on the generation of consensus sequences by *LORCAN*. Uneven spacing of the data points between samples and missing values are a result of differences in the fraction of input reads used to generate the top consensus sequences. **A**) Percent identity of each consensus sequence against the consensus sequences produced from the full dataset. **B**) Numbers of ambiguous bases, **C**) number of gaps, and **D**) total length of the consensus sequences.

**Figure 5.**
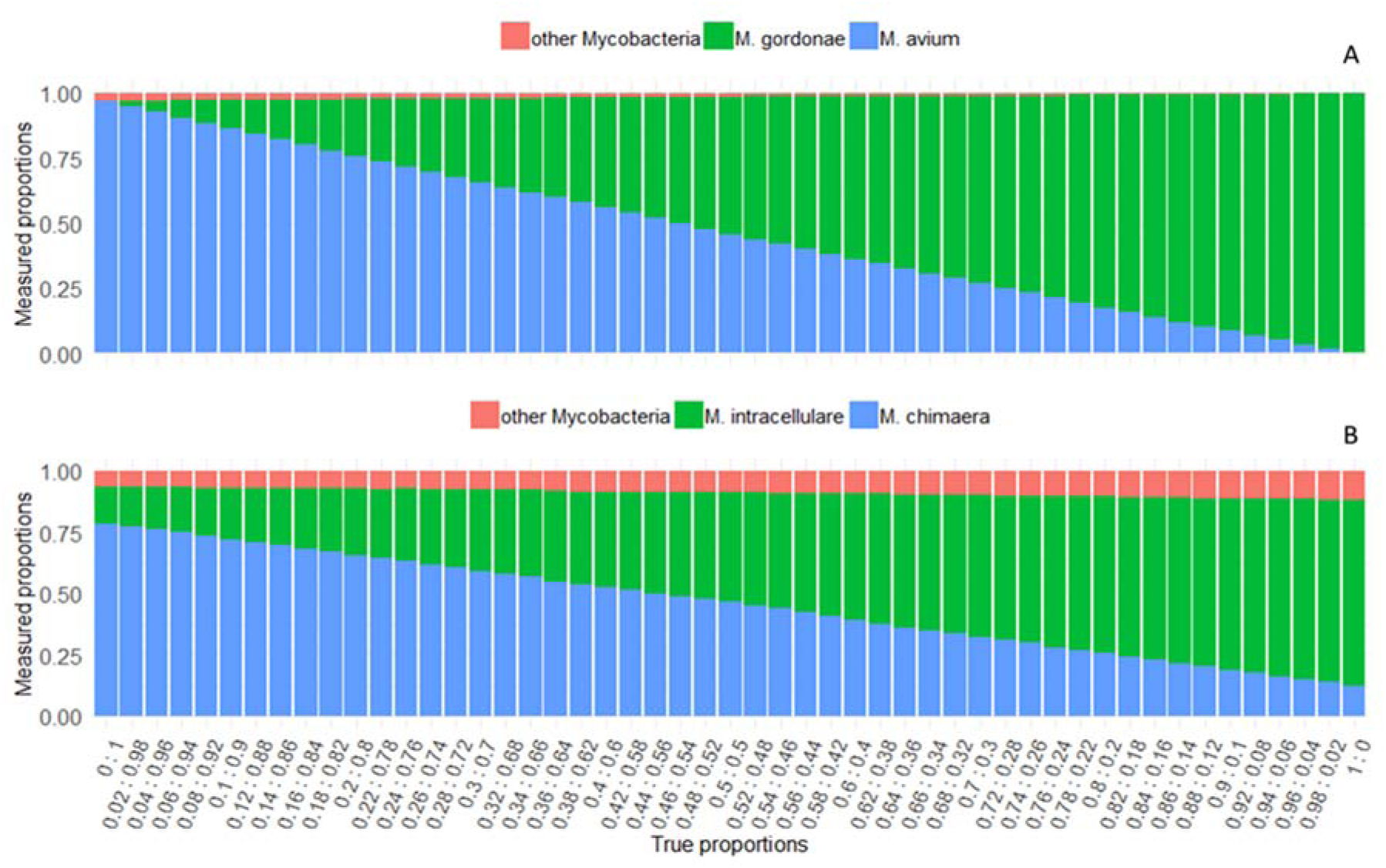
*LORCAN* analysis of artificial read mixtures produced from pairs of *Mycobacterium* species with different genetic relatedness: **A**) *M. gordonae* and *M. avium*, sequence identity: 97.64% (950/973 bases identical). **B**) *M. chimaera* and *M. intracellulare*, sequence identity 99.79% (976/978 bases identical). X-axis represent the true compositions of the read mixtures. Example: x-values of 0.1:0.9 represent mixtures of 10% *M. gordonae* and 90% *M. avium* (**A**) and 10% *M. chimaera* and 90% *M. intracellulare* (**B**).

**Figure 6.**
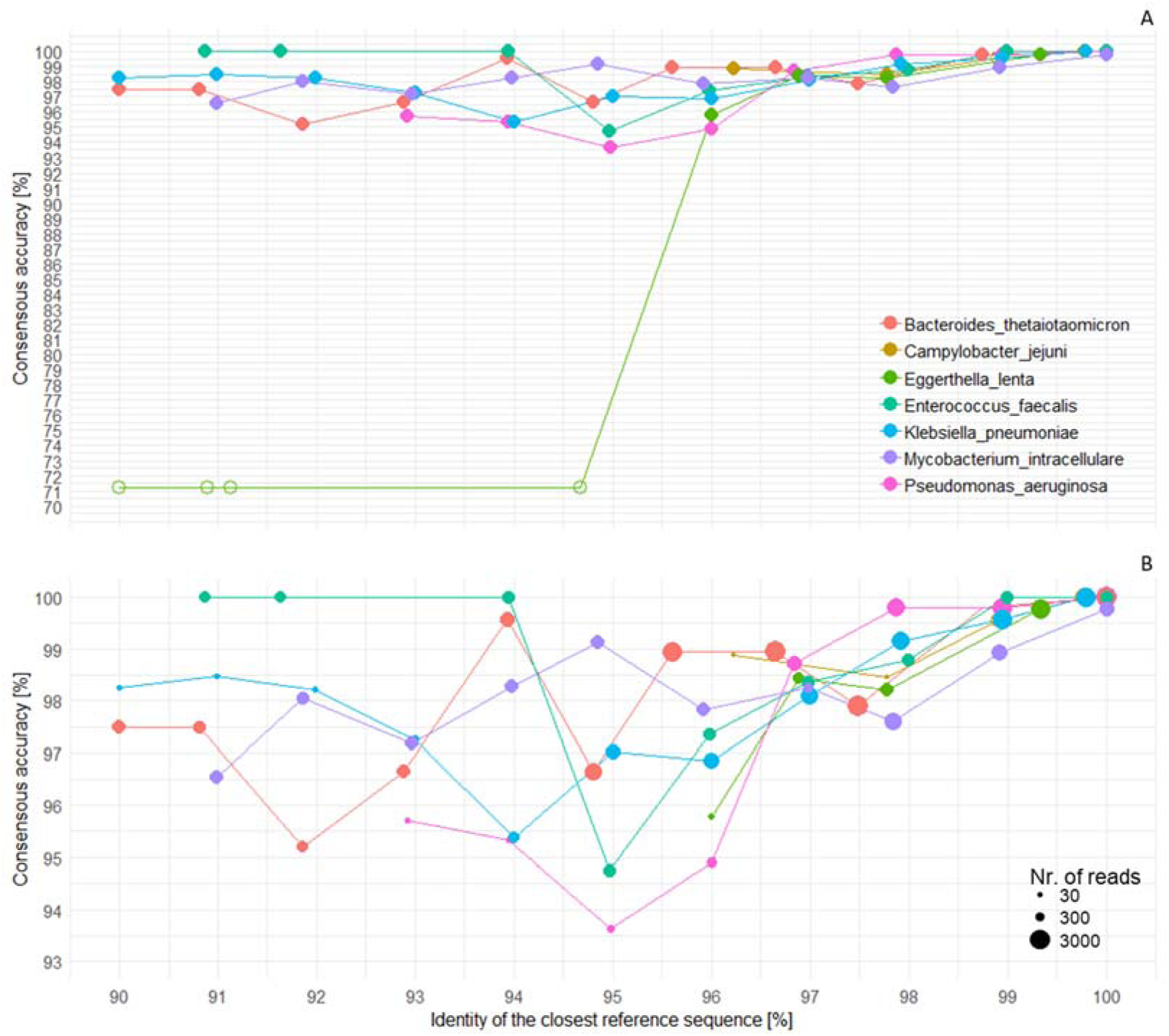
Influence of reference database completeness on consensus sequence generation. Each consensus sequence was compared to a consensus sequence produced with a perfectly matching reference sequence. Additionally, each consensus sequence was identified by *BLAST* similarity search against the full reference database. The uneven spacing of the data points reflects the database composition after subsetting. Missing values are a result of insufficient numbers of reads mapping to the reference database. **A**) Filled circles indicate correct taxonomic identification of the ATCC strains. The low identities and unsuccessful identification of *Eggerthella lenta* are a result of a low-level contamination in combination with unsuccessful mapping of the *Eggerthella* reads. **B**) the diameter of the circles is proportional to the number of reads used for consensus generation. Additional details are provided in Table S2 and Figure S4.

### Data availability

All reads and sequences corresponding to the data presented in Table 1, Figures 4 and 6, were deposited to the European Nucleotide Archive, under the project reference PRJEB34167. Sanger and *LORCAN*-derived consensus sequences are available as supplementary multi-FASTA files.

**Table 1.**
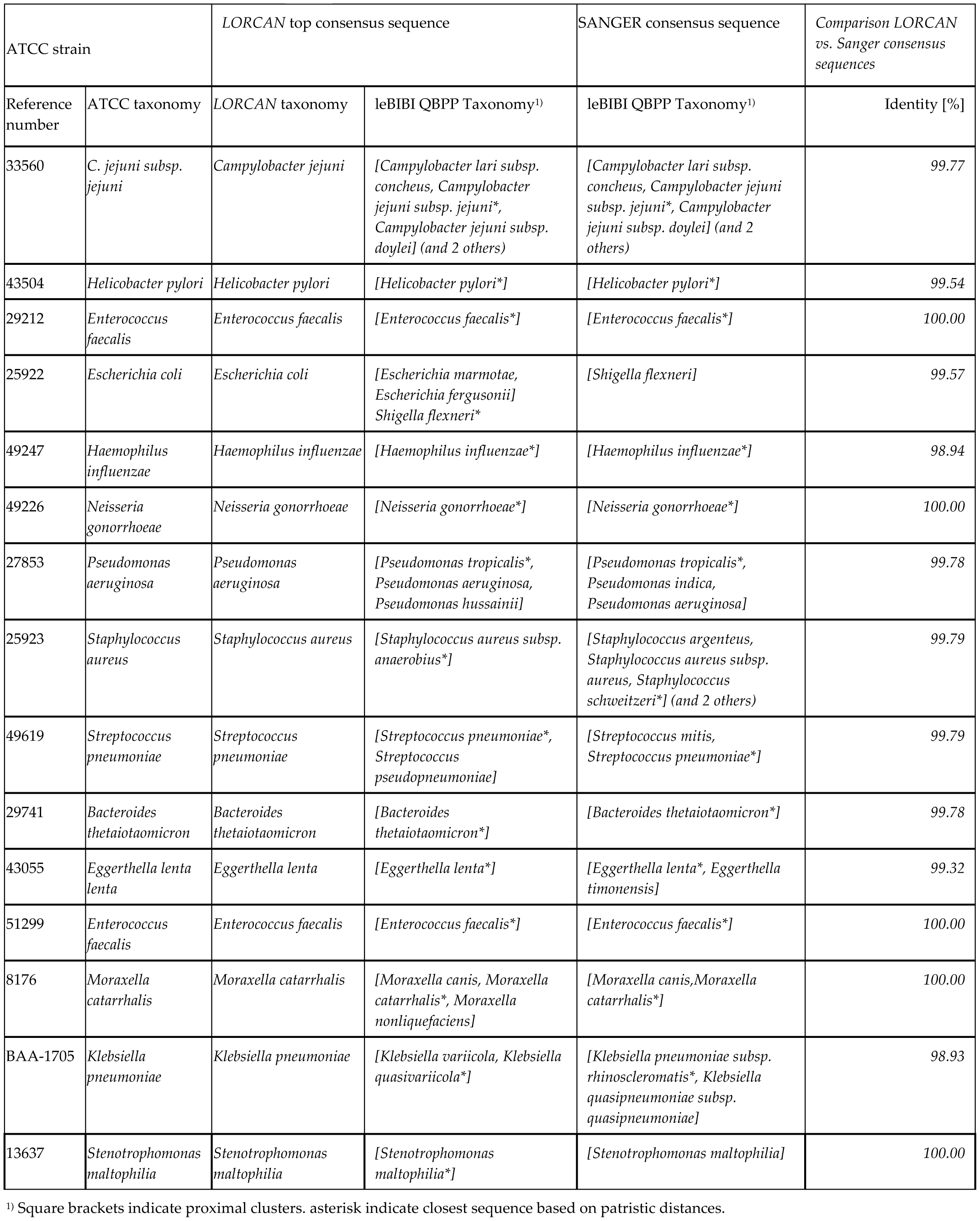
Validation of taxonomic classification of ATCC reference strains. Samples were analysed in parallel by Sanger sequencing and with the approach presented in this study, and the resulting consensus sequences were submitted to the online identification platform leBIBI QBPP.

## Results

We developed a standardized laboratory workflow, as well as the fully automated analysis pipeline, which together provide a sample-to-report solution for taxonomic identification of bacterial cultures based on amplicon sequencing of their 16S rRNA genes. The laboratory workflow, which was tested and adjusted for parallel processing of up to 16 samples done manually by a single person (theoretically scalable up to 96 samples using automation), includes stringent quality control steps to guarantee consistent results. The analysis pipeline is based on publicly available software and runs on Linux machines. It automates quality control, demultiplexing, consensus sequence generation, taxonomic analysis based on the highly curated leBIBI 16S database, as well as report generation (text, PDF). Validation was conducted by direct comparison to Sanger sequencing with real clinical samples consisting of pure or mixed rRNA amplicons belonging to several bacterial genera (*Bacteroides, Eggerthella, Enterococcus, Klebsiella, Mycobacterium, Campylobacter, Pseudomonas*). Additionally, we created artificial read mixtures from closely related bacterial species to assess the pipeline’s performance and robustness when confronted to contaminated samples. We demonstrated that by combining ONT sequencing and *LORCAN*, the accuracy of Sanger sequencing can be closely matched (>99.6% sequence identity on average) and that mixed samples can be resolved at the single base resolution level.

### Parameter evaluation and optimisation

#### Read length vs. consensus quality

Raw amplicon reads sequenced with the ONT technology may vary in length, as they represent the existing heterogeneity in PCR amplicon sizes. Therefore, to test the influence of read length on quality of the generated consensus sequences, different length fractions (window size 10 bases, 50 reads per window) were analysed with *LORCAN*. With increasing read length higher proportions of the analysed read-sets were used for consensus generation, consensus length increased and numbers of ambiguous bases and gaps decreased (Figure 3A-D). As a compromise between minimal read loss and maximal consensus quality, the following size boundaries (relative to the mode of the read length distribution) were used in this study: Lower boundary defined as the modal length minus 20 bases; upper boundary, as the model length plus 100 bases (Figure 3E).

#### Numbers of read vs. consensus quality

To establish the minimal numbers of reads required to produce high quality consensus sequences, we took the reads obtained from sequencing the 16S amplicons from a given ATCC and randomly sampled subsets of reads ranging from 10 to 1000 reads per subset, and created 100 such subsets. This procedure was repeated with six ATCC strains. All datasets were analysed with *LORCAN* and consensus sequence lengths, numbers of ambiguous bases (n’s), gaps, as well as identities to the corresponding consensus sequences produced from full datasets were determined (Figure 4). With the exception of *K. pneumoniae*, which contains eight 16S rRNA operons of varying sequence lengths (Figures S1 and S2), all *LORCAN*-derived consensus sequences showed improvements with increasing numbers of input reads. The most significant improvements took place between 10 and 100 reads. All consensus sequences produced from more than 100 reads showed identities of 99% or higher to the consensus sequences produced from the full read sets.

#### Validation of SNV discrimination and analysis of mixed samples

To test the ability of *LORCAN* to resolve mixed samples, reads from either *M. intracellulare* and *M. chimaera* (identity: 99.79%), or from *M. avium* and *M. gordonae* (identity: 97.64%), were mixed together at various proportions and analysed with *LORCAN. Mycobacteria* strains were chosen because they include many species with near-identical sequences (in the analysed amplicon region) and are therefore suitable to test the limitations of our approach. Sequence mixtures were resolved accurately if sequence similarities were moderate (e.g. 97.6 % identity; Figure 5A). In mixtures of near-identical sequences, moderate levels of crosstalk were detected (e.g. detection of *M. intracellulare* in a pure *M. chimaera* sample; Figure 5B). The dominant species was, however, correctly identified in all cases.

#### Influence of database completeness on consensus accuracy and taxonomic classification

We analysed the influence of reference database completeness on the resulting consensus accuracy and quality by creating incomplete reference databases, from which we excluded reference sequences if they were too close to the ideal reference sequence, and then performed *LORCAN* analysis with each of these truncated databases in turn. The genetic distances of the closest reference sequences in the reference database strongly influenced the accuracy of the resulting consensus sequences. For instance, *Enterococcus faecalis* showed minimal consensus accuracy at ≤ 95% database identity (Figure 6). This was caused by gaps in the closest reference sequence available. At database identities ≤ 94% identity, the reference sequence with the identified gaps was absent and consensus quality increased again (Figures S5 and S6). Classification at the species level was, however, virtually unaffected in pure samples. The *Eggerthella lenta* dataset contained a contamination of *Pseudomonas stutzeri* reads (0.8% of all reads), which did not influence classification when reference sequences that enabled a mapping of *Eggerthella lenta* reads were available. In the absence of sufficiently close reference sequences, the sample was misidentified (Figure 6A). The numbers provided in the *LORCAN* report did, however, reveal that the *Pseudomonas stutzeri* consensus sequence was only based on 20 out of 850 reads, which therefore indicated a likely case of sub-optimal taxonomic classification.

#### *Validation of sequence consensuses generated by the combination of nanopore sequencing and* LORCAN

The comparison of 75 *LORCAN* generated consensus sequences from 14 sequencing runs (including 61 clinical samples and 14 ATCC reference strains) to their corresponding Sanger sequences revealed an average sequence identity of 99.6% ± 0.5 (standard deviation). The positive control (produced from the same pool of amplicons) that was systematically sequenced in these 14 runs showed an average identity of 99.77 % ± 0.23 to its corresponding Sanger sequence. All reference strains were correctly identified at the species level by *LORCAN*. Identification by LeBIBI QBPP resulted in assignment of the expected species (lowest patristic distance) or the placement of the expected species in the proximal cluster of the query sequence (in the phylogenetic tree) in all but two cases. In these cases, the analysed strains were placed in close neighbourhood of the expected species in the phylogenetic tree produced by LeBIBI QBPP (Table 1, Figure S7).

## Discussion

We present here the first sample-to-report solution for marker-gene based taxonomic identification of bacterial cultures specifically designed for clinical applications. We extensively tested the influences of various analysis parameters and therefore provide a basis for optimal tuning of the *LORCAN* pipeline to specific requirements. We demonstrated that reads significantly shorter than the modal read length showed reduced mappability and that resulting consensus sequences were of reduced quality. No such observations were made when using reads from longer length fractions. Therefore, we excluded reads that were significantly shorter than the mode of the read length distribution (by 20 bases) from the analysis with the corresponding command line parameter in *LORCAN*. With these parameters being set, accurate consensus sequences (≥ 99% identity to sanger sequences produced from the same DNA) were reliably produced with as few as 100 size-filtered reads per sample confirming previous findings [42].

The required number of input reads may vary with the taxonomic complexity of the analysed samples and the resolution required by the operator. From a theoretical viewpoint (Figure 1, step 2), a total of 3,000 size-selected reads may allow for the creation of high-quality consensus sequences and reliable species identification for species contributing ≥3.3% of all reads among those 3.000 selected reads (i.e. when setting a minimum reference mapping depth of 100 reads as *LORCAN* parameter, which corresponds to the minimum number recommended of reads for reliable consensus creation; Figure 4). In most cases, however, even when a sample consists of amplicons derived from a unique species, not all reads may be assigned to the target species (e.g. due to read errors and/or the presence of highly similar sequences associated with other species). Furthermore, demultiplexing and size selection result in significant reduction of available reads. As illustrative purposes, during our last 12 sequencing runs consisting of 49 clinical samples, 15 reference strains, 12 positive controls and 12 place-holders, we produced an average of 664,765 ± SD=269,339 basecalled reads while multiplexing on average 7 ± 3 barcoded samples. Read demultiplexing produced thereafter an average of 14,959 ± 15,635 reads (17% of all reads) with correctly assigned barcode sequences. This comparably high read loss resulted from the stringent demultiplexing parameters used (detection of both 3′ and 5′ barcodes required, exclusion of reads with internal barcodes), which effectively prevent crosstalk between libraries [43]. Subsequent size selection (mode read length -20 to +100 bp) resulted in an average of 13,362 ± 13,593 reads per barcode that were available for further processing. Barcodes with more than 3,000 reads in the vicinity of the expected amplicon size were further down-sampled (threshold of 3000 reads, adjustable *LORCAN* parameter), resulting in an average number of reads of 2,716 ± 521 reads per sample. All samples, controls and place-holders processed in these 12 sequencing runs were successfully taxonomically identified. Although species identification could have been achieved with a lower number of reads per sample, sequence production was fast (i.e. approximately 2-3 hours for 1 million reads), and even if flow cells may have been reused up to three times, the maximal sequencing capacity of the flow cells was never utilized (Table S1).

The compatibility with mixed samples was a key requirement during development of *LORCAN* as contaminations are not rare in cultures derived from clinical samples. To exclude external sources of variation we tested this by analysing artificial read mixtures. *LORCAN* showed high robustness against such mixture events and was capable of quantitatively represent read compositions in mixed samples, as long as the involved species did not have near-identical marker gene regions (Figure 5). In the latter case, the individual species were correctly identified and their consensus sequences were of high quality, but their proportions were moderately skewed.

One potential pitfall of reference-based approaches for consensus building may be the dependency on database completeness. We explored this extensively with custom built databases which lacked reference sequences closely related to the analysed strains. Consensus accuracy was strongly affected, and in order to reliably reach sequence qualities on par with the quality obtained with the Sanger method, *LORCAN* required reference databases of high quality and completeness. Even if databases contained sequences with up to 99% identity to the analysed species, further improvements could often be made by adding closer reference sequences (Figure 6). When the consensus sequence was constructed, however, taxonomic identification based on the obtained consensus sequence was far less sensitive to database completeness: Even consensus sequences produced with distant reference sequences (≤ 90% identity to the analysed strains, using an incomplete database) allowed for reliable identification at the species level, when the generated consensus was compared with a complete database. For clinical strains where database quality is generally high, this finding indicates a high reliability despite the database dependency. This was confirmed by extensive validation in our diagnostics department, which was based on the dual sequencing of clinical samples in parallel with Sanger sequencing and *LORCAN* over several months, which overall showed average sequence identities of 99.6% (and 99.77% for positive controls sequenced conjointly with the clinical samples).

A number of studies on ONT-based marker gene analysis have been published over the past years, covering a range of different laboratory and computational approaches aiming to obtain high quality sequences from ONT reads. Most computational workflows ether include reference-based consensus generation or *de novo* assembly, in combination with additional error correction steps. They were reported to perform similarly in terms of the accuracy of the produced sequences [22,23,25,27,42]. *De novo* approaches are preferable when reference sequences are missing, however, so far the only studies demonstrating “reference-free” consensus generation from complex samples (e.g. mock communities) relied on rather laborious wet-lab procedures such as rolling cycle amplification or unique tagging of the individual amplicons before sequencing [25,26]. Unlike previous studies we specifically designed our workflow for clinical routine applications. Compatibility with mixed samples and time/cost efficacy were therefore key requirements and comprehensive reference databases were readily available. We therefore chose a reference-based approach allowing us to separate reads originating from mixed cultures while using standard ONT protocols. Furthermore, and in contrast to most previous studies, we omitted consensus error correction which is commonly applied to remove homopolymer errors from consensus sequences and assemblies produced from nanopore reads [22,23] because we did not detect a negative influence of the latter errors in our taxonomic classification approach.

The strengths of the nanopore-*LORCAN* approach is that overall the procedure is faster, more flexible, and more cost effective than Sanger or Illumina-based approach, as it relies on both straightforward ONT protocols and automated sample analysis up to result reporting. In addition, nanopore sequencing is compatible with any amplicon size, which is a clear advantage over other existing sequencing technologies, and also allows the processing and resolution of mixed amplicon sample as demonstrated here. Finally, even when the reference sequence database is incomplete or lacks closely related reference sequences, we showed that the approach is robust and provides correct taxonomic identification of the bacterial species. The approach has, however, some limitations, including the limited taxonomic resolution inherent to single-gene based methods. Commonly used 16S rRNA gene regions for example have been reported to allow for genus identification in >90% of cases, for species identification in 65 to 83% of cases and to result in unsuccessful identification in 1 to 14% of all analysed isolates [18,44,45]. Limitations specific to *LORCAN* include that it is currently database-dependent and that high accuracy consensus sequences may require not only complete databases, but also identification of the genetic region of interest. The overall wet laboratory procedure still takes several hours, and would need to be optimized to allow fast and efficient processing of several samples via automation or via simplified steps.

In conclusion, we demonstrate that the combination of nanopore sequencing and *LORCAN* pipeline offers a significant improvement over the well-established Sanger sequencing-based approach, in terms of reliability, flexibility, turnover time, and reproducibility of the results. The described workflow was successfully introduced in the routine of our diagnostics department and has the potential to significantly facilitate amplicon sequencing in other diagnostic settings.

## Acknowledgements

We thank Heiko Amlang and Christian Baumann for their excellent technical assistance, and John W Looney for his assistance in the preparation of technical documents.

## List of Supplementary Documents

### Supporting Figures and Tables

**Figure S1**. *LORCAN* consensus sequence aligned to *K. pneumoniae* reference sequences.

**Figure S2.** *LORCAN K. pneumoniae* consensus plot.

**Figure S3**. Pairwise alignments of *M. avium, M. gordonae, M. intracellulare and M. chimaera.*

**Figure S4.** The influence of the similarity of database completeness on consensus sequence quality.

**Figure S5.** Alignment of *Enterococcu*s consensus sequences with the sequence used for their generation.

**Figure S6**. *Enterococcus faecalis* consensus sequences together with the 10 best *BLAST* hits.

**Table S1.** Number of reads at different steps of the *LORCAN* pipeline.

**Table S2.** The influence of the similarity of database completeness on consensus sequence quality.

**Table S3.** Third-party software utilised in the *LORCAN* pipeline.

**<u>Supplementary Text S1</u>.** Detailed wet-laboratory protocol.

**<u>LORCAN_reports.tar.gz</u>**. Reports of the *LORCAN* pipeline including as .PDF and .TXT files (ATCC strains in Table 1).

**<u>LORCAN_consensus.fa</u>.** Multi-FASTA file of the *LORCAN*-generated consensus sequences for the ATCC strains (Table 1).

**<u>SANGER_consensus.fa</u>.** Multi-FASTA file of the Sanger sequences for the ATCC strains (Table 1).

## References

1. Keys C.J., Dare D.J., Sutton H., Wells G., Lunt M., McKenna T., McDowall M., Shah H.N. Compilation of a MALDI-TOF mass spectral database for the rapid screening and characterisation of bacteria implicated in human infectious diseases. Infect Genet Evol 2004, 4, pages:221–42.

2. Barberis C., Almuzara M., Join-Lambert O., Ramirez M.S., Famiglietti A., Vay C. Comparison of the Bruker MALDI-TOF mass spectrometry system and conventional phenotypic methods for identification of Gram-positive rods. PLoS One 2014, 9, pages:e106303.

3. Levesque S., Dufresne P.J., Soualhine H., Domingo M.C., Bekal S., Lefebvre B., Tremblay C. A Side by Side Comparison of Bruker Biotyper and VITEK MS: Utility of MALDI-TOF MS Technology for Microorganism Identification in a Public Health Reference Laboratory. PLoS One 2015, 10, pages:e0144878.

4. Bizzini A., Jaton K., Romo D., Bille J., Prod’hom G., Greub G. Matrix-assisted laser desorption ionization-time of flight mass spectrometry as an alternative to 16S rRNA gene sequencing for identification of difficult-to-identify bacterial strains. J Clin Microbiol 2011, 49, pages:693–6.

5. Rocca M.F., Barrios R., Zintgraff J., Martinez C., Irazu L., Vay C., Prieto M. Utility of platforms Viteks MS and Microflex LT for the identification of complex clinical isolates that require molecular methods for their taxonomic classification. PLoS One 2019, 14, pages:e0218077.

6. Sandalakis V., Goniotakis I., Vranakis I., Chochlakis D., Psaroulaki A. Use of MALDI-TOF mass spectrometry in the battle against bacterial infectious diseases: recent achievements and future perspectives. Expert Rev Proteomics 2017, 14, pages:253–67.

7. Psaroulaki A., Chochlakis D. Use of MALDI-TOF mass spectrometry in the battle against bacterial infectious diseases: recent achievements and future perspectives. Expert Rev Proteomics 2018, 15, pages:537–39.

8. Tsuchida S. Application of MALDI-TOF for Bacterial Identification. The Use of Mass Spectrometry Technology (MALDI-TOF) in Clinical Microbiology,. 2018, pages:101–12.

9. Ramulu H.G., Groussin M., Talla E., Planel R., Daubin V., Brochier-Armanet C. Ribosomal proteins: toward a next generation standard for prokaryotic systematics? Mol Phylogenet Evol 2014, 75, pages:103–17.

10. Jauffrit F., Penel S., Delmotte S., Rey C., de Vienne D.M., Gouy M., Charrier J.P., Flandrois J.P., Brochier-Armanet C. RiboDB Database: A Comprehensive Resource for Prokaryotic Systematics. Mol Biol Evol 2016, 33, pages:2170–2.

11. Maruvada P., Leone V., Kaplan L.M., Chang E.B. The Human Microbiome and Obesity: Moving beyond Associations. Cell Host Microbe 2017, 22, pages:589–99.

12. Durban A., Abellan J.J., Jimenez-Hernandez N., Ponce M., Ponce J., Sala T., D’Auria G., Latorre A., Moya A. Assessing gut microbial diversity from feces and rectal mucosa. Microb Ecol 2011, 61, pages:123–33.

13. Janda J.M., Abbott S.L. 16S rRNA gene sequencing for bacterial identification in the diagnostic laboratory: pluses, perils, and pitfalls. J Clin Microbiol 2007, 45, pages:2761–4.

14. Srinivasan R., Karaoz U., Volegova M., MacKichan J., Kato-Maeda M., Miller S., Nadarajan R., Brodie E.L., Lynch S.V. Use of 16S rRNA gene for identification of a broad range of clinically relevant bacterial pathogens. PLoS One 2015, 10, pages:e0117617.

15. Zhang J., Fang Y., Hou J.Y., Ren H.J., Jiang R., Roos P., Dovichi N.J. Use of non-cross-linked polyacrylamide for four-color DNA sequencing by capillary electrophoresis separation of fragments up to 640 bases in length in two hours. Anal Chem 1995, 67, pages:4589–93.

16. Heather J.M., Chain B. The sequence of sequencers: The history of sequencing DNA. Genomics 2016, 107, pages:1–8.

17. Tenney A.E., Wu J.Q., Langton L., Klueh P., Quatrano R., Brent M.R. A tale of two templates: automatically resolving double traces has many applications, including efficient PCR-based elucidation of alternative splices. Genome Res 2007, 17, pages:212–8.

18. Mignard S., Flandrois J.P. 16S rRNA sequencing in routine bacterial identification: a 30-month experiment. J Microbiol Methods 2006, 67, pages:574–81.

19. Nicholls S.M., Quick J.C., Tang S., Loman N.J. Ultra-deep, long-read nanopore sequencing of mock microbial community standards. Gigascience 2019, 8, pages.

20. Srivathsan A., Baloglu B., Wang W., Tan W.X., Bertrand D., Ng A.H.Q., Boey E.J.H., Koh J.J.Y., Nagarajan N., Meier R. A MinION-based pipeline for fast and cost-effective DNA barcoding. Mol Ecol Resour 2018, pages.

21. Vaser R., Sovic I., Nagarajan N., Sikic M. Fast and accurate de novo genome assembly from long uncorrected reads. Genome Res 2017, 27, pages:737–46.

22. Menegon M., Cantaloni C., Rodriguez-Prieto A., Centomo C., Abdelfattah A., Rossato M., Bernardi M., Xumerle L., Loader S., Delledonne M. On site DNA barcoding by nanopore sequencing. PLoS One 2017, 12, pages:e0184741.

23. Maestri S., Cosentino E., Paterno M., Freitag H., Garces J.M., Marcolungo L., Alfano M., Njunjic I., Schilthuizen M., Slik F., Menegon M., Rossato M., Delledonne M. A Rapid and Accurate MinION-Based Workflow for Tracking Species Biodiversity in the Field. Genes (Basel) 2019, 10, pages.

24. Somerville V., Lutz S., Schmid M., Frei D., Moser A., Irmler S., Frey J.E., Ahrens C.H. Long-read based de novo assembly of low-complexity metagenome samples results in finished genomes and reveals insights into strain diversity and an active phage system. BMC Microbiol 2019, 19, pages:143.

25. Calus S.T., Ijaz U.Z., Pinto A.J. NanoAmpli-Seq: a workflow for amplicon sequencing for mixed microbial communities on the nanopore sequencing platform. Gigascience 2018, 7, pages.

26. Karst S.M., Ziels R.M., Kirkegaard R.H., Albertsen M. Enabling high-accuracy long-read amplicon sequences using unique molecular identifiers and Nanopore sequencing. bioRxiv 2019, pages:645903.

27. Benitez-Paez A., Portune K.J., Sanz Y. Species-level resolution of 16S rRNA gene amplicons sequenced through the MinION portable nanopore sequencer. Gigascience 2016, 5, pages:4.

28. Kai S., Matsuo Y., Nakagawa S., Kryukov K., Matsukawa S., Tanaka H., Iwai T., Imanishi T., Hirota K. Rapid bacterial identification by direct PCR amplification of 16S rRNA genes using the MinION nanopore sequencer. FEBS Open Bio 2019, 9, pages:548–57.

29. Ramette A. GitHub Repository for LORCAN Pipeline. Available online: https://github.com/aramette/LORCAN/

30. Wick R.R. Available online: https://github.com/rrwick/Porechop. pages.

31. Li H. Minimap2: pairwise alignment for nucleotide sequences. Bioinformatics 2018, 34, pages:3094–100.

32. Li H., Handsaker B., Wysoker A., Fennell T., Ruan J., Homer N., Marth G., Abecasis G., Durbin R., Genome Project Data Processing S. The Sequence Alignment/Map format and SAMtools. Bioinformatics 2009, 25, pages:2078–9.

33. Shen W., Le S., Li Y., Hu F. SeqKit: A Cross-Platform and Ultrafast Toolkit for FASTA/Q File Manipulation. PLoS One 2016, 11, pages:e0163962.

34. Altschul S.F., Gish W., Miller W., Myers E.W., Lipman D.J. Basic local alignment search tool. J Mol Biol 1990, 215, pages:403–10.

35. Katoh K., Standley D.M. MAFFT multiple sequence alignment software version 7: improvements in performance and usability. Mol Biol Evol 2013, 30, pages:772–80.

36. Castresana J. Selection of conserved blocks from multiple alignments for their use in phylogenetic analysis. Mol Biol Evol 2000, 17, pages:540–52.

37. Nguyen L.T., Schmidt H.A., von Haeseler A., Minh B.Q. IQ-TREE: a fast and effective stochastic algorithm for estimating maximum-likelihood phylogenies. Mol Biol Evol 2015, 32, pages:268–74.

38. Flandrois J.P., Perriere G., Gouy M. leBIBIQBPP: a set of databases and a webtool for automatic phylogenetic analysis of prokaryotic sequences. BMC Bioinformatics 2015, 16, pages:251.

39. Schloss P.D., Westcott S.L., Ryabin T., Hall J.R., Hartmann M., Hollister E.B., Lesniewski R.A., Oakley B.B., Parks D.H., Robinson C.J., Sahl J.W., Stres B., Thallinger G.G., Van Horn D.J., Weber C.F. Introducing mothur: open-source, platform-independent, community-supported software for describing and comparing microbial communities. Appl Environ Microbiol 2009, 75, pages:7537–41.

40. leBIBI-QBPP. https://umr5558-bibiserv.univ-lyon1.fr/lebibi/lebibi.cgi, database procaryota_SSU-rDNA-16S_TS-stringent, version 2019/Feb/07 14:40.

41. Corpet F. Multiple sequence alignment with hierarchical clustering. Nucleic Acids Res 1988, 16, pages:10881–90.

42. Pomerantz A., Penafiel N., Arteaga A., Bustamante L., Pichardo F., Coloma L.A., Barrio-Amoros C.L., Salazar-Valenzuela D., Prost S. Real-time DNA barcoding in a rainforest using nanopore sequencing: opportunities for rapid biodiversity assessments and local capacity building. Gigascience 2018, 7, pages.

43. Xu Y., Lewandowski K., Lumley S., Pullan S., Vipond R., Carroll M., Foster D., Matthews P.C., Peto T., Crook D. Detection of Viral Pathogens With Multiplex Nanopore MinION Sequencing: Be Careful With Cross-Talk. Front Microbiol 2018, 9, pages:2225–25.

44. Woo P.C., Ng K.H., Lau S.K., Yip K.T., Fung A.M., Leung K.W., Tam D.M., Que T.L., Yuen K.Y. Usefulness of the MicroSeq 500 16S ribosomal DNA-based bacterial identification system for identification of clinically significant bacterial isolates with ambiguous biochemical profiles. J Clin Microbiol 2003, 41, pages:1996–2001.

45. Drancourt M., Bollet C., Carlioz A., Martelin R., Gayral J.P., Raoult D. 16S ribosomal DNA sequence analysis of a large collection of environmental and clinical unidentifiable bacterial isolates. J Clin Microbiol 2000, 38, pages:3623–30.

