## Supplementary material for "A sample-to-report solution for taxonomic identification of cultured bacteria in the clinical setting based on nanopore sequencing": Supporting Figures and Tables.pdf

### Supporting materials

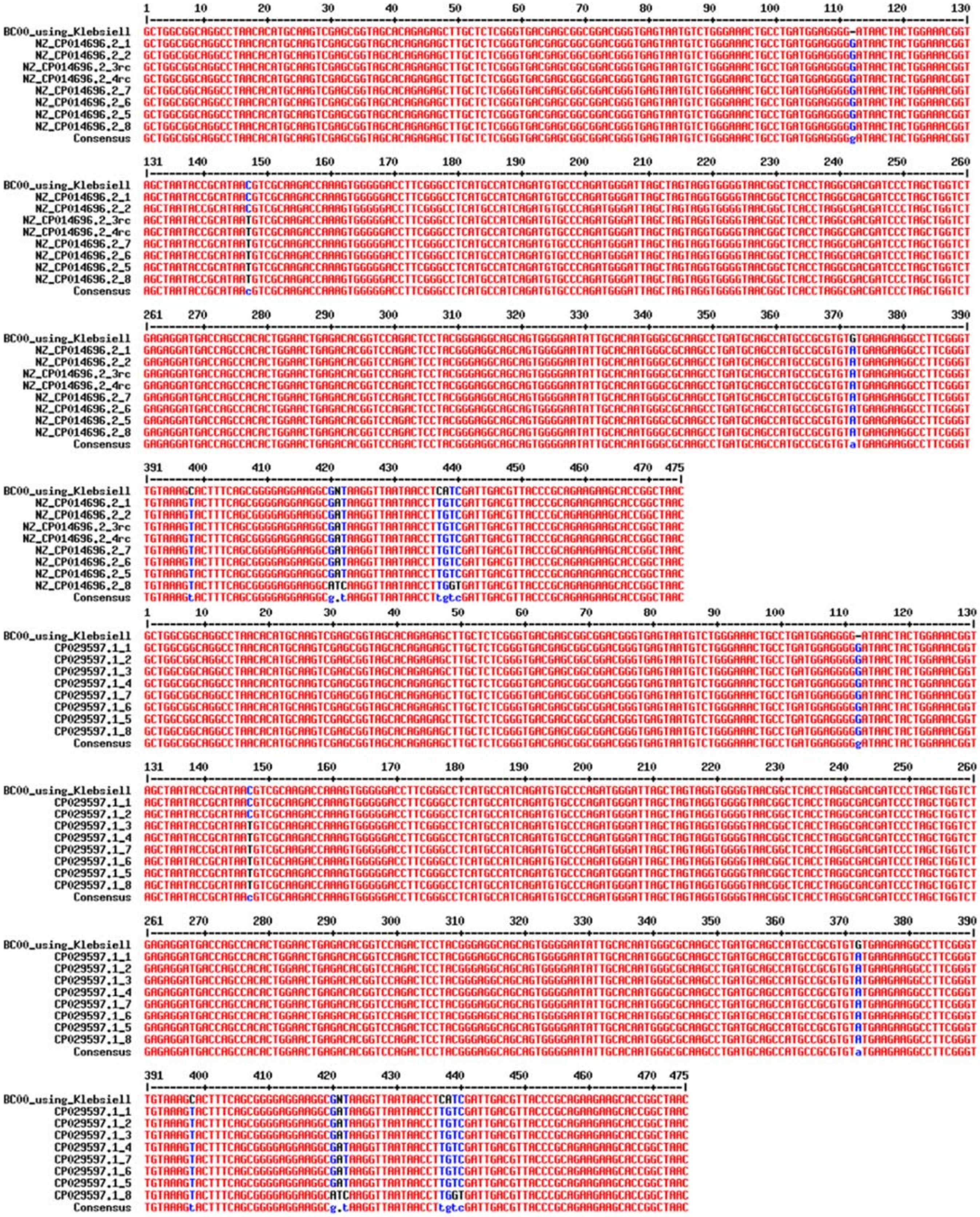

**Figure S1.** *K. pneumoniae* LORCAN consensus (prefix "BC00") aligned to the corresponding regions of the 8 rRNA operons of two published genomes of the same ATCC reference strain.

### Figure S1

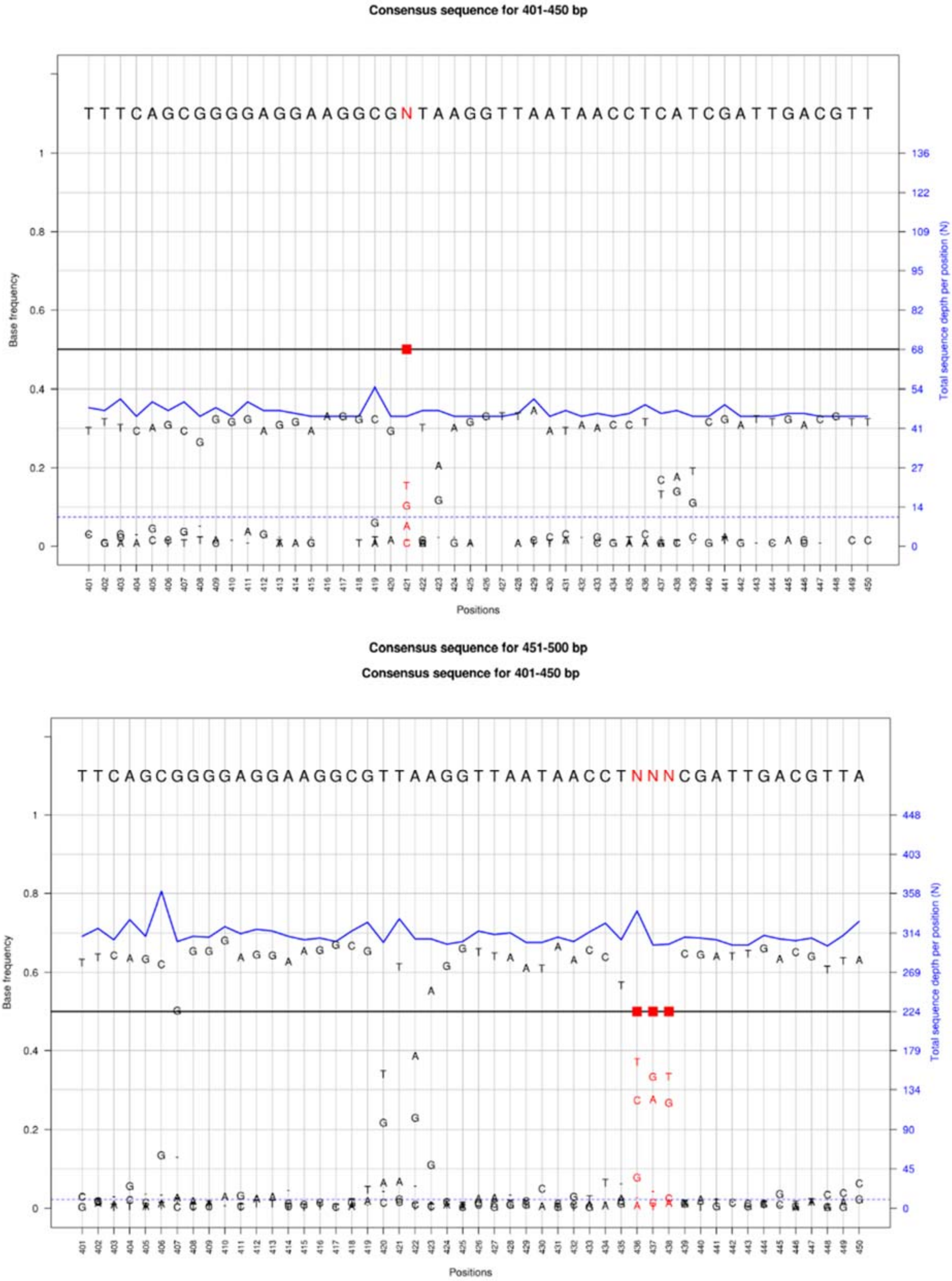


**Figure S2.** *K. pneumoniae* consensus plots (*LORCAN* output) of consensus sequences produced from different numbers of reads randomly selected from the same dataset. Upper panel: 600 reads, bottom panel: 300 reads. N's are located in positions where the operons of the *K. pneumoniae* strain show variability.

#### Figure S2

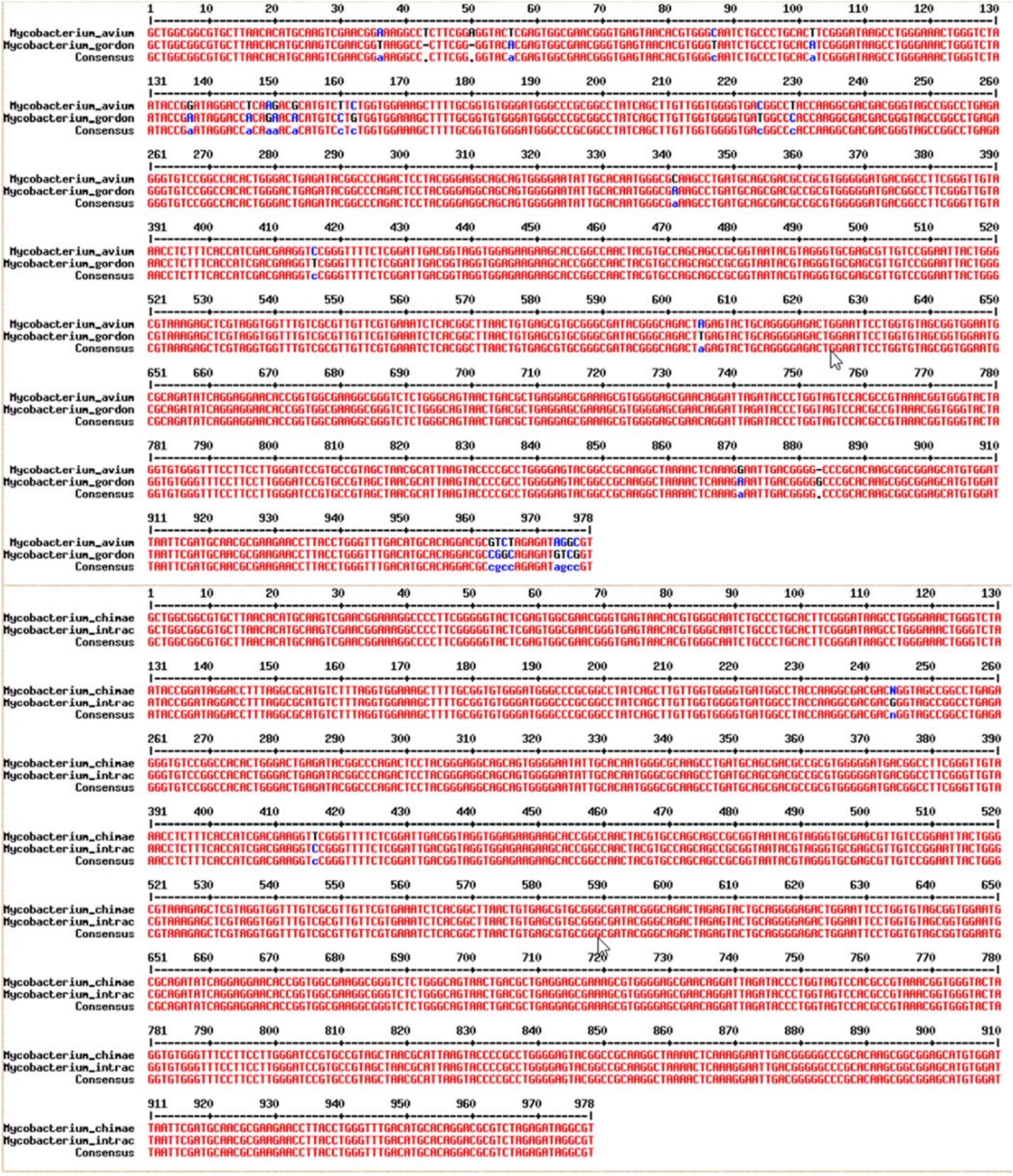

**Figure S3.** Pairwise alignments of *M. avium and M. gordonae* (upper panel)*, M. intracellulare* and *M. chimaera* (bottom panel) 16S rRNA gene regions used for accessing the performance of *LORCAN* with mixed samples.

#### Figure S3

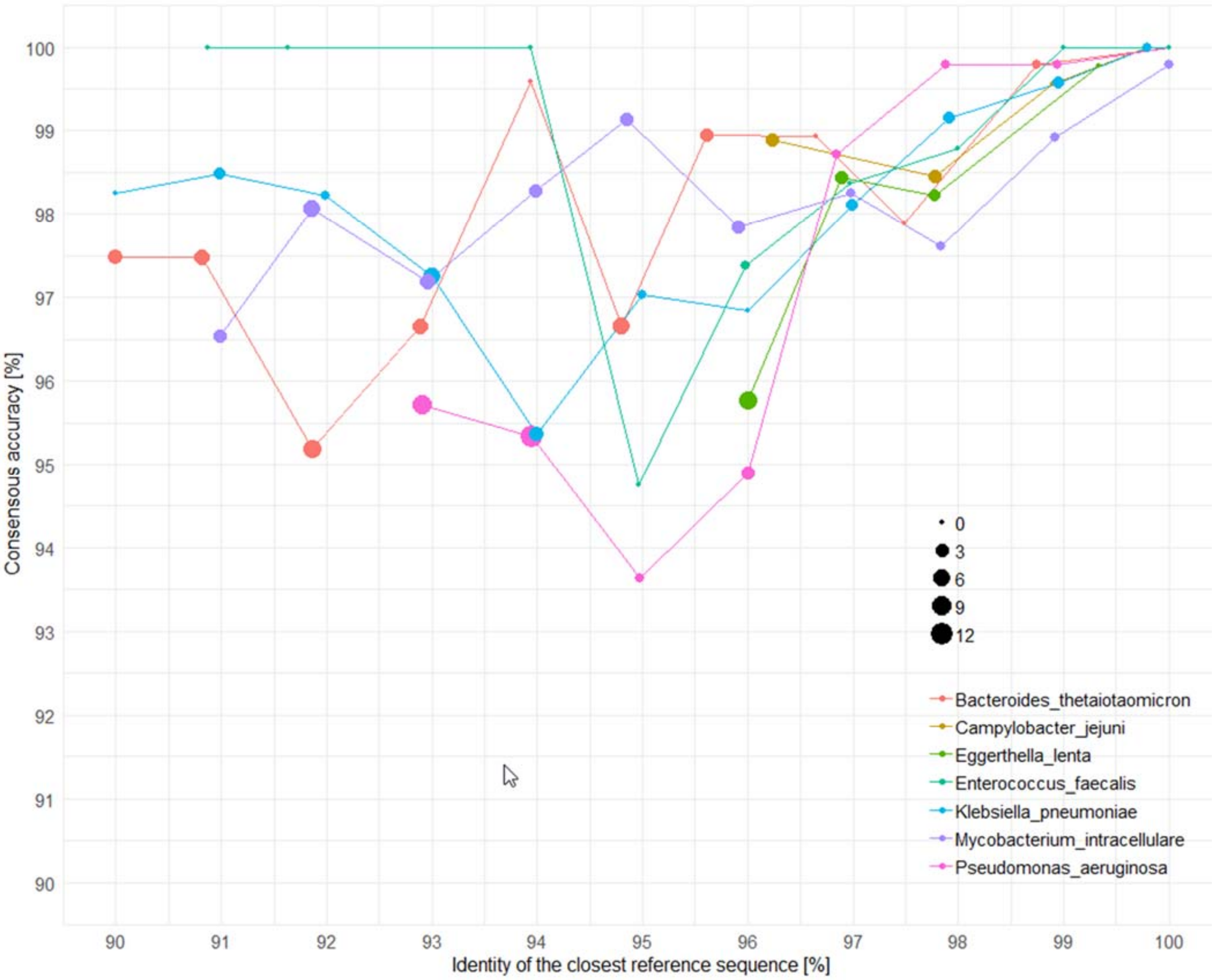


**Figure S4**. Influence of the similarity of the analysed reads with their best match in the reference database on consensus accuracy. Each consensus sequence was compared to a consensus sequence produced with a perfectly matching reference sequence. The sizes of the dots represent numbers of "N" in the consensus sequence. Missing points are a result of insufficient numbers of reads mapping to the reference database.

#### Figure S4

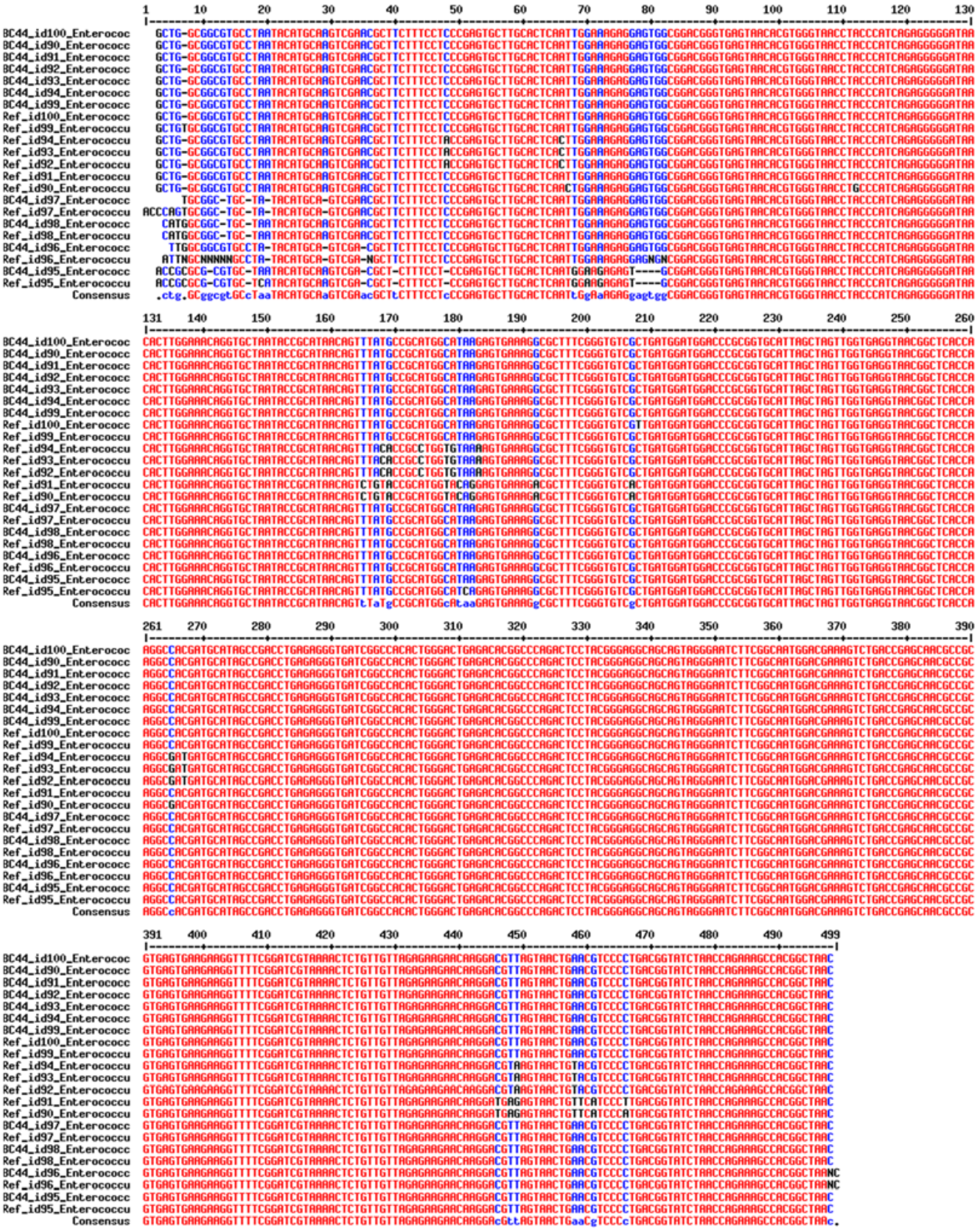

**Figure S5.** Alignment of *Enterococcus* consensus sequences with the sequence used for their generation. Reference sequences are marked with the prefix "Ref", *LORCAN* consensus sequences with the prefix "BC44, and the number following "id" indicates the identity threshold used to subset the database before consensus generation. Example: "BC44_id95_: Consensus sequence generated with a database where all references with sequence identities of ≥ 95% to the analysed strain were removed. "Ref_id95": the closest remaining reference sequence after subsetting of the database.

### Figure S5

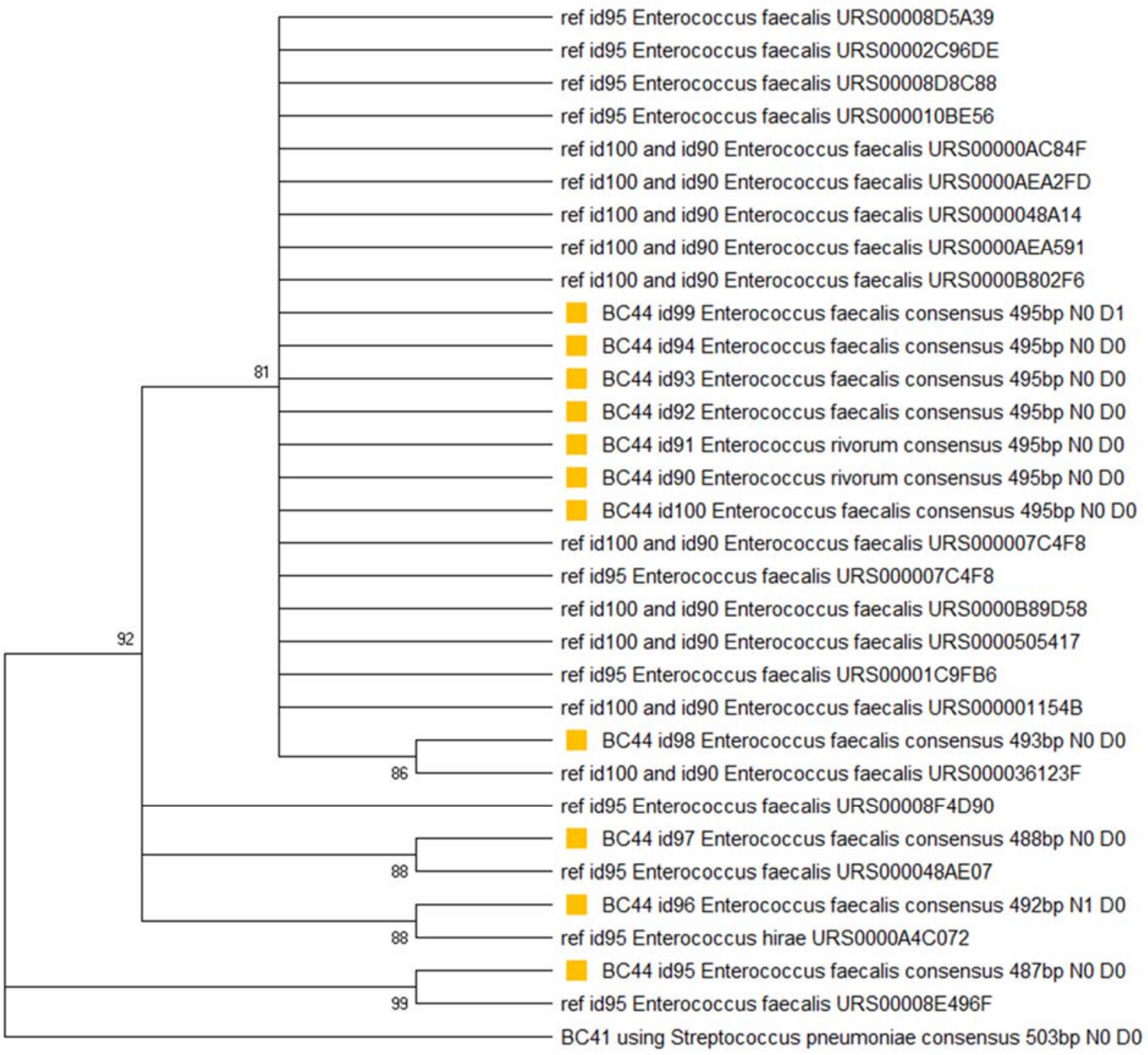


**Figure S6.** Evolutionary relationships of consensus sequences and their reference sequences. *Enterococcus faecalis* consensus (marked by yellow boxes) sequences together with the 10 best BLAST-hits of the consensus sequences produced with a 100%, a 95% and a 90% identical reference sequence (marked by the prefixes id100, id95 and id90). The evolutionary history was inferred using the UPGMA method. The bootstrap consensus tree inferred from 1000 replicates is taken to represent the evolutionary history of the taxa analysed. Branches corresponding to partitions reproduced in less than 80% bootstrap replicates are collapsed. The percentage of replicate trees in which the associated taxa clustered together in the bootstrap test (1000 replicates) are shown next to the branches. The evolutionary distances were computed using the Maximum Composite Likelihood method and are in the units of the number of base substitutions per site. This analysis involved 32 nucleotide sequences. All ambiguous positions were removed for each sequence pair (pairwise deletion option). There were a total of 521 positions in the final dataset. Evolutionary analyses were conducted in MEGA X software.

#### Figure S6

A

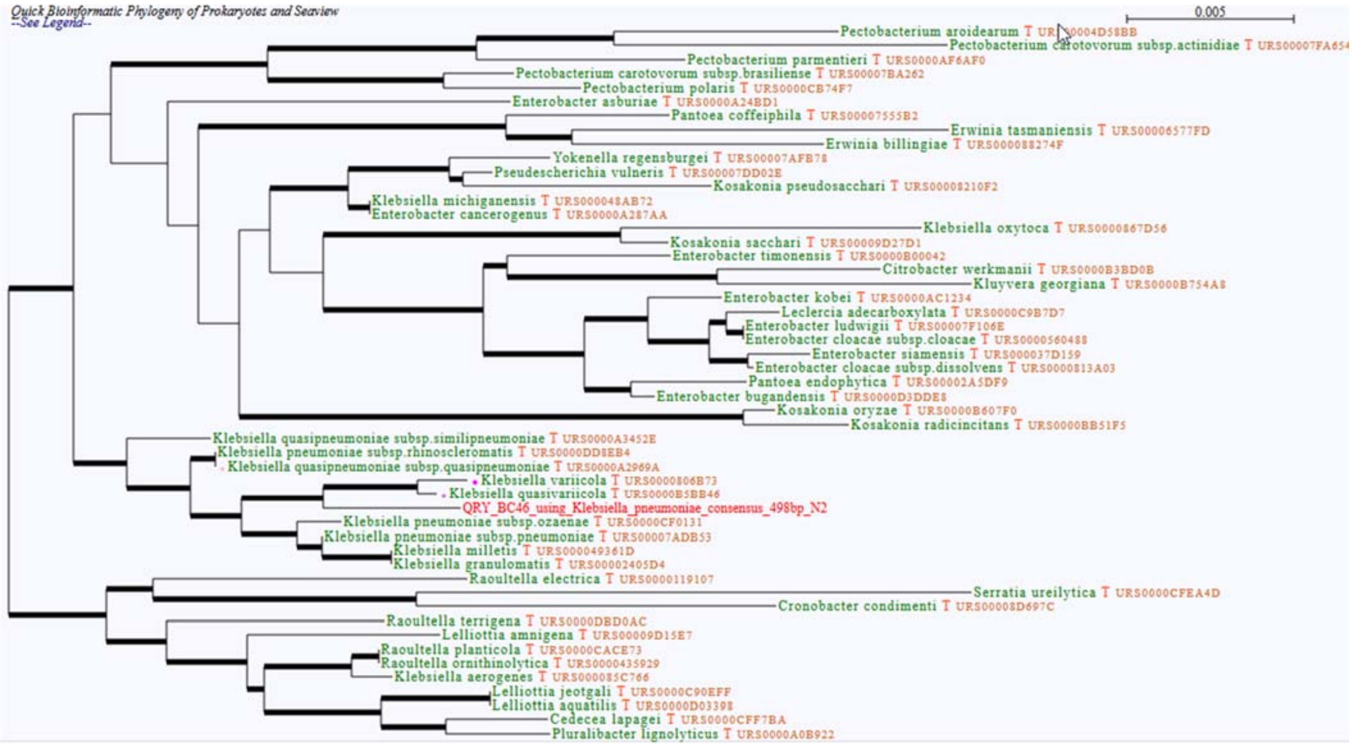
Quick Bioinformatic Phylogeny of Prokaryotes and Seaview
--See Legend--
0.005
Pectobacterium aroidearum T URS0004D58BB
Pectobacterium carotovorum subsp.actinidiae T URS00007FA654
Pectobacterium parmentieri T URS0000AF6AF0
Pectobacterium carotovorum subsp.brasiliense T URS00007BA262
Pectobacterium polaris T URS0000CB74F7
Enterobacter asburiae T URS0000A24BD1
Pantoea coffeiphila T URS00007555B2
Erwinia tasmaniensis T URS00006577FD
Erwinia billingiae T URS000088274F
Yokenella regensburgei T URS00007AFB78
Pseudescherichia vulneris T URS00007DD02E
Kosakonia pseudosacchari T URS00008210F2
Klebsiella michiganensis T URS000048AB72
Enterobacter cancerogenus T URS0000A287AA
Klebsiella oxytoca T URS0000867D56
Kosakonia sacchari T URS00009D27D1
Enterobacter timonensis T URS0000B00042
Citrobacter werkmanii T URS0000B3BD0B
Kluyvera georgiana T URS0000B754A8
Enterobacter kobei T URS0000AC1234
Leclercia adecarboxylata T URS0000C9B7D7
Enterobacter ludwigii T URS00007F106E
Enterobacter cloacae subsp.cloacae T URS0000560488
Enterobacter siamensis T URS000037D159
Enterobacter cloacae subsp.dissolvens T URS0000813A03
Pantoea endophytica T URS00002A5DF9
Enterobacter bugandensis T URS0000D3DDE8
Kosakonia oryzae T URS0000B607F0
Kosakonia radicincitans T URS0000BB51F5
Klebsiella quasipneumoniae subsp.similipneumoniae T URS0000A3452E
Klebsiella pneumoniae subsp.rhinoscleromatis T URS0000DD8EB4
Klebsiella quasipneumoniae subsp.quasipneumoniae T URS0000A2969A
Klebsiella variicola T URS0000806B73
Klebsiella quasivariicola T URS0000B5BB46
QRY_BC46_using_Klebsiella_pneumoniae_consensus_498bp_N2
Klebsiella pneumoniae subsp.ozaenae T URS0000CF0131
Klebsiella pneumoniae subsp.pneumoniae T URS00007ADB53
Klebsiella milletis T URS000049361D
Klebsiella granulomatis T URS00002405D4
Raoultella electrica T URS0000119107
Serratia ureilytica T URS0000CFEA4D
Cronobacter condimenti T URS00008D697C
Raoultella terrigena T URS0000DBD0AC
Lelliottia amnigena T URS00009D15E7
Raoultella planticola T URS0000CACE73
Raoultella ornithinolytica T URS0000435929
Klebsiella aerogenes T URS000085C766
Lelliottia jeotgali T URS0000C90EFF
Lelliottia aquatilis T URS0000D03398
Cedecea lapagei T URS0000CFF7BA
Pluralibacter lignolyticus T URS0000A0B922

B

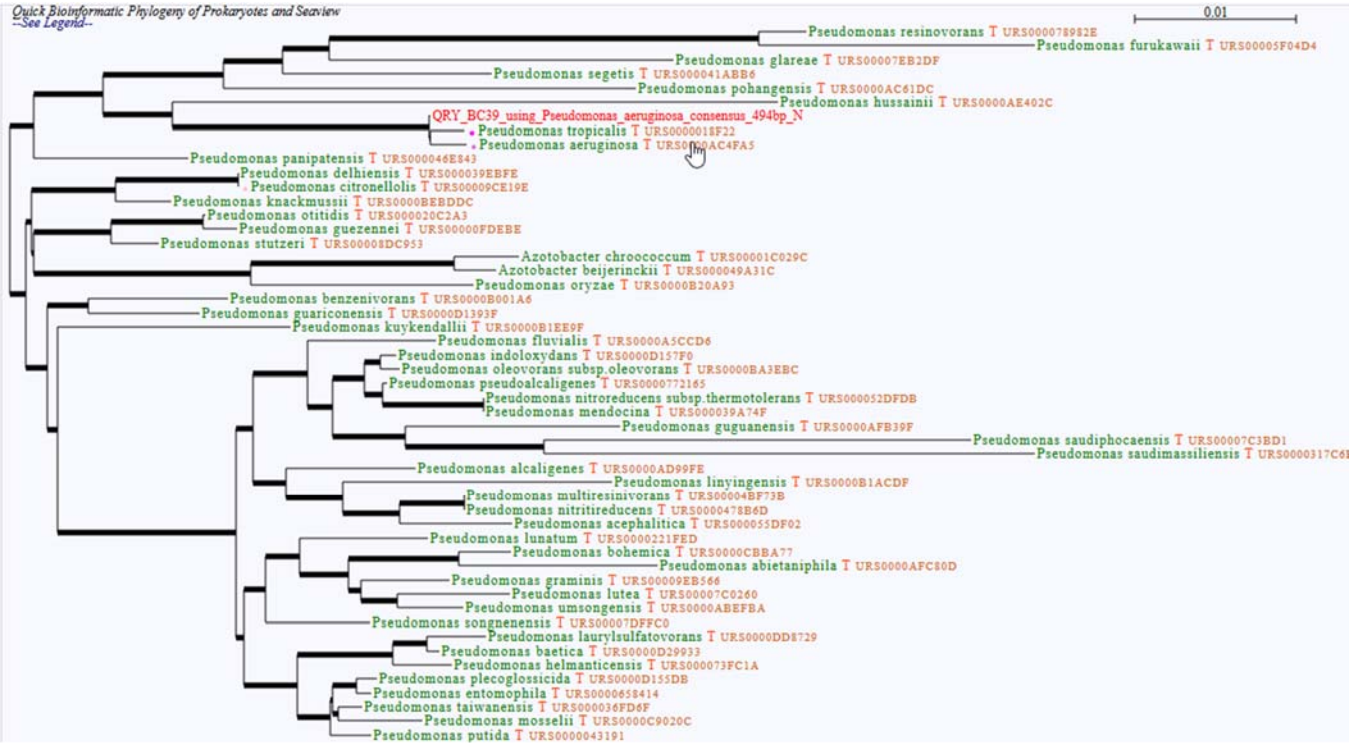
Quick Bioinformatic Phylogeny of Prokaryotes and Seaview
--See Legend--
0.01
Pseudomonas resinovorans T URS000078982E
Pseudomonas furukawaii T URS00005F04D4
Pseudomonas glareae T URS00007EB2DF
Pseudomonas segetis T URS000041ABB6
Pseudomonas pohangensis T URS0000AC61DC
Pseudomonas hussainii T URS0000AE402C
QRY_BC39_using_Pseudomonas_aeruginosa_consensus_494bp_N
Pseudomonas tropicalis T URS000018F22
Pseudomonas aeruginosa T URS0000AC4FA5
Pseudomonas panipatensis T URS000046E843
Pseudomonas delhiensis T URS000039EBFE
Pseudomonas citronellolis T URS00009CE19E
Pseudomonas knackmussii T URS0000BEBDDC
Pseudomonas otitidis T URS000020C2A3
Pseudomonas guezennei T URS00000FDEBE
Pseudomonas stutzeri T URS00008DC953
Azotobacter chroococcum T URS00001C029C
Azotobacter beijerinckii T URS000049A31C
Pseudomonas oryzae T URS0000B20A93
Pseudomonas benzenivorans T URS0000B001A6
Pseudomonas guariconensis T URS0000D1393F
Pseudomonas kuykendallii T URS0000B1EE9F
Pseudomonas fluvialis T URS0000A5CCD6
Pseudomonas indoloxydans T URS0000D157F0
Pseudomonas oleovorans subsp.oleovorans T URS0000BA3EBC
Pseudomonas pseudoalcaligenes T URS0000772165
Pseudomonas nitroreducens subsp.thermotolerans T URS000052DFDB
Pseudomonas mendocina T URS000039A74F
Pseudomonas guguanensis T URS0000AFB39F
Pseudomonas saudiphocaensis T URS00007C3BD1
Pseudomonas saudimassiliensis T URS0000317C61
Pseudomonas alcaligenes T URS0000AD99FE
Pseudomonas linyingensis T URS0000B1ACDF
Pseudomonas multiresinivorans T URS00004BF73B
Pseudomonas nitritireducens T URS0000478B6D
Pseudomonas acephalitica T URS000055DF02
Pseudomonas lunatum T URS0000221FED
Pseudomonas bohemica T URS0000CBBA77
Pseudomonas abietaniphila T URS0000AFC80D
Pseudomonas graminis T URS00009EB566
Pseudomonas lutea T URS00007C0260
Pseudomonas umsongensis T URS0000ABEFBA
Pseudomonas songnenensis T URS00007DFFC0
Pseudomonas laurylsulfatovorans T URS0000DD8729
Pseudomonas baetica T URS0000D29933
Pseudomonas helmanticensis T URS000073FC1A
Pseudomonas plecoglossicida T URS0000D155DB
Pseudomonas entomophila T URS0000658414
Pseudomonas taiwanensis T URS000036FD6F
Pseudomonas mosselii T URS0000C9020C
Pseudomonas putida T URS0000043191

C

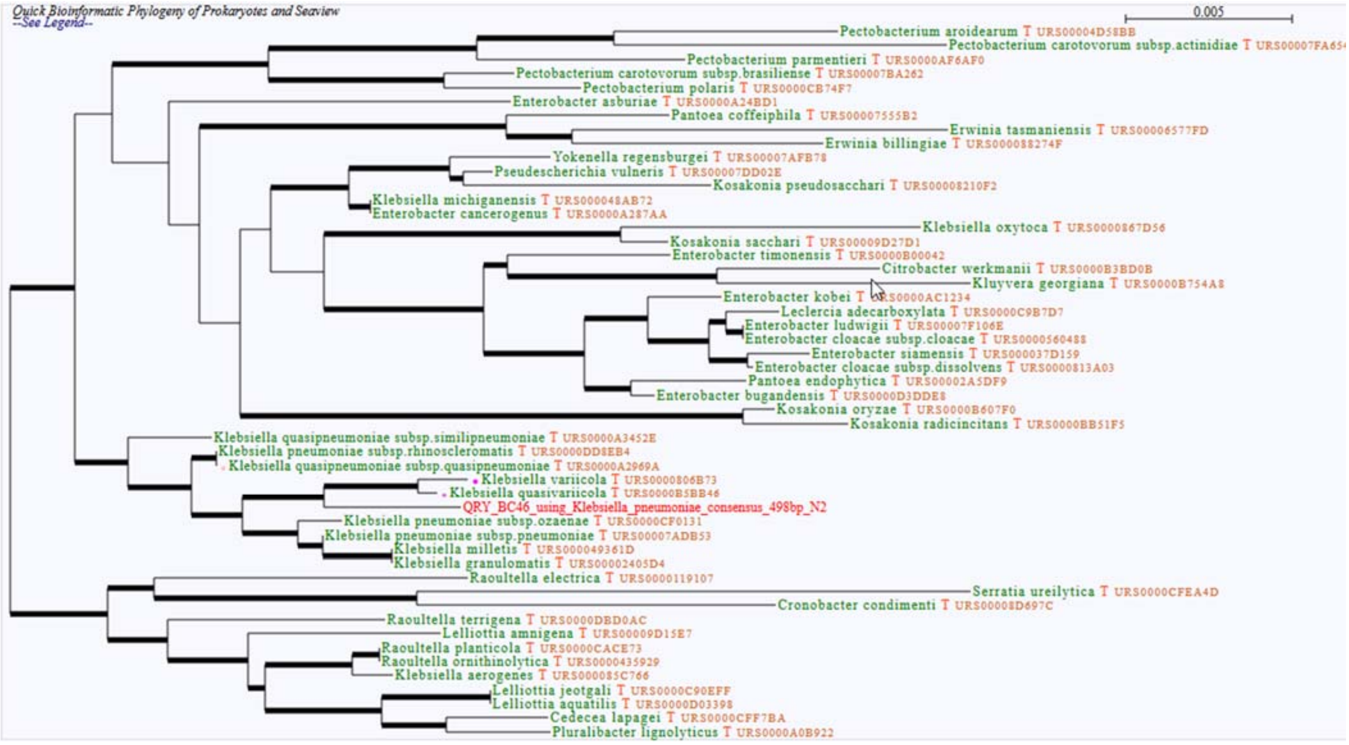
Quick Bioinformatic Phylogeny of Prokaryotes and Seaview
--See Legend--
0.005
Pectobacterium aroidearum T URS00004D58BB
Pectobacterium carotovorum subsp.actinidiae T URS00007FA65
Pectobacterium parmentieri T URS0000AF6AF0
Pectobacterium carotovorum subsp.brasiliense T URS00007BA262
Pectobacterium polaris T URS0000CB74F7
Enterobacter asburiae T URS0000A24BD1
Pantoea coffeiphila T URS00007555B2
Erwinia tasmaniensis T URS00006577FD
Erwinia billingiae T URS000088274F
Yokenella regensburgei T URS00007AFB78
Pseudescherichia vulneris T URS00007DD02E
Kosakonia pseudosacchari T URS00008210F2
Klebsiella michiganensis T URS000048AB72
Enterobacter cancerogenus T URS0000A287AA
Klebsiella oxytoca T URS0000867D56
Kosakonia sacchari T URS00009D27D1
Enterobacter timonensis T URS0000B00042
Citrobacter werkmanii T URS0000B3BD0B
Kluyvera georgiana T URS0000B754A8
Enterobacter kobei T URS0000AC1234
Leclercia adecarboxylata T URS0000C9B7D7
Enterobacter ludwigii T URS00007F106E
Enterobacter cloacae subsp.cloacae T URS0000560488
Enterobacter siamensis T URS000037D159
Enterobacter cloacae subsp.dissolvens T URS0000813A03
Pantoea endophytica T URS00002A5DF9
Enterobacter bugandensis T URS0000D3DDE8
Kosakonia oryzae T URS0000B607F0
Kosakonia radicincitans T URS0000BB51F5
Klebsiella quasipneumoniae subsp.similipneumoniae T URS0000A3452E
Klebsiella pneumoniae subsp.rhinoscleromatis T URS0000DD8EB4
Klebsiella quasipneumoniae subsp.quasipneumoniae T URS0000A2969A
Klebsiella variicola T URS0000806B73
Klebsiella quasivariicola T URS0000B5BB46
QRY_BC46_using_Klebsiella_pneumoniae_consensus_498bp_N2
Klebsiella pneumoniae subsp.ozaenae T URS0000CF0131
Klebsiella pneumoniae subsp.pneumoniae T URS00007ADB53
Klebsiella milletis T URS000049361D
Klebsiella granulomatis T URS00002405D4
Raoultella electrica T URS0000119107
Serratia ureilytica T URS0000CFEA4D
Cronobacter condimenti T URS00008D697C
Raoultella terrigena T URS0000DBD0AC
Lelliottia amnigena T URS00009D15E7
Raoultella planticola T URS0000CACE73
Raoultella ornithinolytica T URS0000435929
Klebsiella aerogenes T URS000085C766
Lelliottia jeotgali T URS0000C90EFF
Lelliottia aquatilis T URS0000D03398
Cedecea lapagei T URS0000CFF7BA
Pluralibacter lignolyticus T URS0000A0B922

D

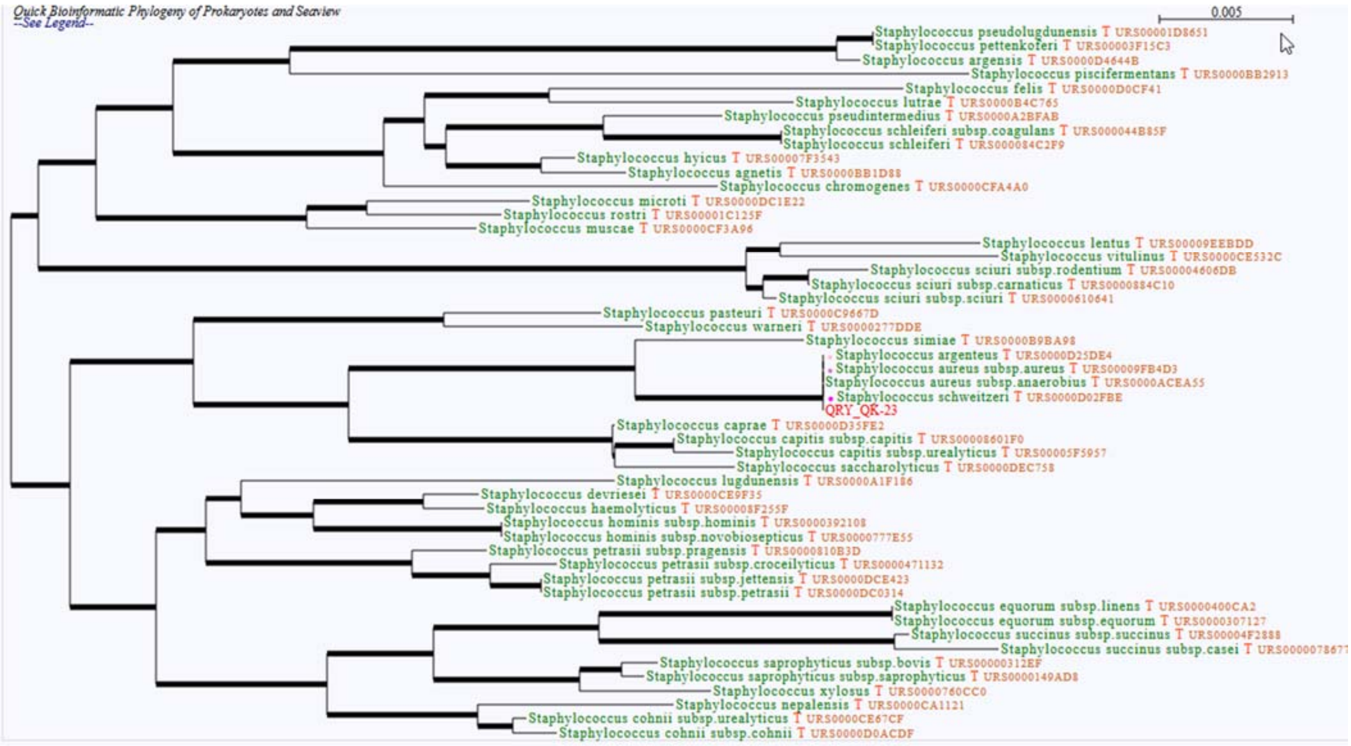
Quick Bioinformatic Phylogeny of Prokaryotes and Seaview
--See Legend--
0.005
Staphylococcus pseudolugdunensis T URS00001D8651
Staphylococcus pettenkoferi T URS00003F15C3
Staphylococcus argensis T URS0000D4644B
Staphylococcus piscifermentans T URS0000BB2913
Staphylococcus felis T URS0000D0CF41
Staphylococcus lutrae T URS0000B4C765
Staphylococcus pseudintermedius T URS0000A2BFAB
Staphylococcus schleiferi subsp.coagulans T URS000044B85F
Staphylococcus schleiferi T URS000084C2F9
Staphylococcus hyicus T URS00007F3543
Staphylococcus agnetis T URS0000BB1D88
Staphylococcus chromogenes T URS0000CFA4A0
Staphylococcus microti T URS0000DC1E22
Staphylococcus rostri T URS00001C125F
Staphylococcus muscae T URS0000CF3A96
Staphylococcus lentus T URS00009EEBDD
Staphylococcus vitulinus T URS0000CE532C
Staphylococcus sciuri subsp.rodentium T URS00004606DB
Staphylococcus sciuri subsp.carnaticus T URS0000884C10
Staphylococcus sciuri subsp.sciuri T URS0000610641
Staphylococcus pasteuri T URS0000C9667D
Staphylococcus warneri T URS0000277DDE
Staphylococcus simiae T URS0000B9BA98
Staphylococcus argenteus T URS0000D25DE4
Staphylococcus aureus subsp.aureus T URS00009FB4D3
Staphylococcus aureus subsp.anaerobius T URS0000ACEA55
Staphylococcus schweitzeri T URS0000D02FBE
QRY_QK-23
Staphylococcus caprae T URS0000D35FE2
Staphylococcus capitis subsp.capitis T URS00008601F0
Staphylococcus capitis subsp.urealyticus T URS00005F5957
Staphylococcus saccharolyticus T URS0000DEC758
Staphylococcus lugdunensis T URS0000A1F186
Staphylococcus devriesei T URS0000CE9F35
Staphylococcus haemolyticus T URS00008F255F
Staphylococcus hominis subsp.hominis T URS0000392108
Staphylococcus hominis subsp.novobiosepticus T URS0000777E55
Staphylococcus petrasii subsp.pragensis T URS0000810B3D
Staphylococcus petrasii subsp.croceilyticus T URS0000471132
Staphylococcus petrasii subsp.jettensis T URS0000DCE423
Staphylococcus petrasii subsp.petrasii T URS0000DC0314
Staphylococcus equorum subsp.linens T URS0000400CA2
Staphylococcus equorum subsp.equorum T URS0000307127
Staphylococcus succinus subsp.succinus T URS00004F2888
Staphylococcus succinus subsp.casei T URS0000078677
Staphylococcus saprophyticus subsp.bovis T URS000000312EF
Staphylococcus saprophyticus subsp.saprophyticus T URS0000149AD8
Staphylococcus xylosus T URS0000760CC0
Staphylococcus nepalensis T URS0000CA1121
Staphylococcus cohnii subsp.urealyticus T URS0000CE67CF
Staphylococcus cohnii subsp.cohnii T URS0000D0ACDF

E

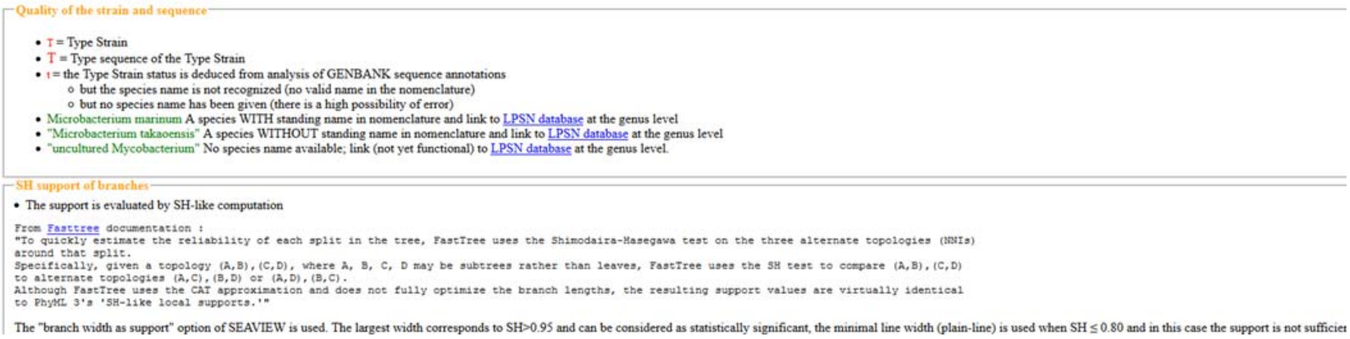

Quality of the strain and sequence

- T = Type Strain
- T = Type sequence of the Type Strain
- t = the Type Strain status is deduced from analysis of GENBANK sequence annotations
    - but the species name is not recognized (no valid name in the nomenclature)
    - but no species name has been given (there is a high possibility of error)
- Microbacterium marinum A species WITH standing name in nomenclature and link to LPSN database at the genus level
- "Microbacterium takaoensis" A species WITHOUT standing name in nomenclature and link to LPSN database at the genus level
- "uncultured Mycobacterium" No species name available; link (not yet functional) to LPSN database at the genus level.

SH support of branches

- The support is evaluated by SH-like computation

```
From Fasttree documentation :
"To quickly estimate the reliability of each split in the tree, FastTree uses the Shimodaira-Hasegawa test on the three alternate topologies (NNIs)
around that split.
Specifically, given a topology (A,B),(C,D), where A, B, C, D may be subtrees rather than leaves, FastTree uses the SH test to compare (A,B),(C,D)
to alternate topologies (A,C),(B,D) or (A,D),(B,C).
Although FastTree uses the CAT approximation and does not fully optimize the branch lengths, the resulting support values are virtually identical
to PhyML 3's 'SH-like local supports.'"
```

The "branch width as support" option of SEAVIEW is used. The largest width corresponds to SH>0.95 and can be considered as statistically significant, the minimal line width (plain-line) is used when SH ≤ 0.80 and in this case the support is not sufficier

**Figure S7.** Phylogenetic trees of strains with ambiguous identification (see main text, Table 1) copied from the leBIBI QBPP outputs. All analysed strains are labelled with the prefix "QRY_". **A**): *Escherichia coli* ATTC 25922 (LORCAN consensus), **B**) *Pseudomonas aeruginosa* ATTC 27853 (LORCAN consensus), **C**) *Klebsiella pneumoniae* ATCC BAA-1705 (LORCAN consensus), **D**) *Staphylococcus aureus* ATCC 25923 (Sanger consensus). **E**) legend copied from the leBIBI QBPP output.

### Figure S7

#### Table S1

**Table S1**: Number of reads at different steps of the *LORCAN* analysis pipeline.

| Nr. of raw basecalled reads | Nr. of samples | Nr. of reads per sample after demultiplexing | | Nr. of reads after size selection | | Nr. of reads after application of the 3000 read cutoff | | Percent of reads used for top consensus generation | |
|---|---|---|---|---|---|---|---|---|---|
| Total | Total | Av. per sample | SD | Av. per sample | SD | Av. per sample | SD | Av. per sample | SD |
| 613032 | 6 | 12043 | 7639 | 13378 | 8506 | 3000.83 | 1.94 | 83.89 | 15.09 |
| 578552 | 7 | 7917 | 2925 | 9597 | 3654 | 3001.14 | 1.21 | 81.49 | 10.55 |
| 221153 | 4 | 10621 | 6798 | 11300 | 7275 | 2990.50 | 15.02 | 59.29 | 19.36 |
| 856179 | 9 | 11708 | 6318 | 12804 | 6452 | 2999.44 | 3.32 | 81.70 | 23.24 |
| 609449 | 6 | 6447 | 7235 | 6990 | 7800 | 1643.00 | 1422.48 | 79.97 | 13.61 |
| 264000 | 7 | 5091 | 3750 | 6097 | 5407 | 2540.14 | 977.25 | 81.26 | 18.41 |
| 784261 | 7 | 14675 | 6973 | 15845 | 7254 | 2990.43 | 8.70 | 64.61 | 13.77 |
| 623687 | 8 | 9331 | 2618 | 10167 | 2664 | 2967.43 | 7.74 | 71.24 | 28.61 |
| 578776 | 9 | 7524 | 3253 | 8642 | 4272 | 2948.11 | 65.86 | 66.58 | 17.25 |
| 937805 | 2 | 55330 | 48012 | 63464 | 58131 | 1630.00 | 1931.82 | 84.18 | 20.77 |
| 1204359 | 7 | 14076 | 3535 | 15280 | 3086 | 2999.86 | 3.24 | 85.78 | 16.32 |
| 705931 | 16 | 5576 | 1995 | 5942 | 2129 | 2882.25 | 472.87 | 94.27 | 9.28 |
| Av: 664765<br>SD: 269339 | Av: 7<br>SD: 3 | Av: 14959<br>SD: 15635 | | Av:.13362<br>SD: 13593 | | Av: 2716<br>SD: 521 | | Av: 78<br>SD: 10 | |

### Table S2

**Table S2:** The influence of the similarity of database completeness on consensus sequence quality: detailed results.

| Sample | Identity cutoff database preparation | Identity of the closest database member to the sample | Identity of the LORCAN consensus sequence to consensus sequences produced from full databases | Numbers of reads mapped | Percent of reads mapped to the top consensus sequences | BLAST identification |
|---|---|---|---|---|---|---|
| *Bacteroides thetaiotaomicron* | 100 | 100.00 | 100.00 | 2092 | 99.95 | correct |
| | 99 | 98.74 | 99.79 | NA | NA | correct |
| | 98 | 97.49 | 97.89 | 2979 | 99.33 | correct |
| | 97 | 96.65 | 98.94 | 2974 | 99.17 | correct |
| | 96 | 95.61 | 98.95 | 2742 | 91.43 | correct |
| | 95 | 94.80 | 96.65 | 1850 | 61.69 | correct |
| | 94 | 93.93 | 99.58 | 1362 | 45.42 | correct |
| | 93 | 92.89 | 96.65 | 1123 | 37.45 | correct |
| | 92 | 91.86 | 95.19 | 604 | 20.14 | correct |
| | 91 | 90.81 | 97.49 | 843 | 28.10 | correct |
| | 90 | 90.00 | 97.49 | 946 | 31.52 | correct |
| *Eggerthella lenta* | 100 | 99.33 | 99.78 | 2530 | 98.75 | correct |
| | 99 | 97.78 | 98.22 | 985 | 96.95 | correct |
| | 98 | 97.78 | 98.22 | 985 | 96.95 | correct |
| | 97 | 96.89 | 98.44 | 520 | 94.20 | correct |
| | 96 | 96.00 | 95.77 | 87 | 69.60 | correct |
| | 95 | 94.67 | 71.21 | 20 | 48.78 | incorrect |
| | 94 | 91.13 | 71.21 | 20 | 55.56 | incorrect |
| | 93 | 91.13 | 71.21 | 20 | 55.56 | incorrect |
| | 92 | 91.13 | 71.21 | 20 | 55.56 | incorrect |
| | 91 | 90.89 | 71.21 | 20 | 55.56 | incorrect |
| | 90 | 90.00 | 71.21 | 20 | 55.56 | incorrect |
| *Enterococcus faecalis* | 100 | 99.78 | 100.00 | 2103 | 73.51 | correct |
| | 100 | 100.00 | 100.00 | 794 | 99.75 | correct |
| | 99 | 98.89 | 99.56 | 984 | 67.91 | incorrect |

| | | | | | | |
|---|---|---|---|---|---|---|
| | 99 | 98.99 | 100.00 | 846 | 99.65 | correct |
| | 98 | 97.78 | 98.45 | 14 | 32.56 | correct |
| | 98 | 97.99 | 98.79 | 837 | 98.59 | correct |
| | 97 | 96.23 | 98.89 | 15 | 78.95 | correct |
| | 97 | 96.98 | 98.36 | 838 | 98.59 | correct |
| | 96 | 95.98 | 97.37 | 830 | 97.76 | correct |
| | 95 | 94.97 | 94.75 | 779 | 91.65 | correct |
| | 94 | 93.94 | 100.00 | 692 | 81.41 | correct |
| | 93 | 91.63 | 100.00 | 590 | 69.41 | correct |
| | 92 | 91.63 | 100.00 | 593 | 69.76 | correct |
| | 91 | 90.87 | 100.00 | 632 | 74.35 | correct |
| | 90 | 89.90 | 100.00 | 357 | 42.00 | correct |
| *Klebsiella pneumoniae* | 100 | 99.79 | 100.00 | 2680 | 90.54 | correct |
| | 99 | 98.95 | 99.58 | 2663 | 90.67 | correct |
| | 98 | 97.92 | 99.16 | 2160 | 78.52 | correct |
| | 97 | 97.00 | 98.11 | 1812 | 68.09 | correct |
| | 96 | 96.00 | 96.84 | 1407 | 62.15 | correct |
| | 95 | 95.00 | 97.03 | 1242 | 80.08 | correct |
| | 94 | 94.00 | 95.37 | 592 | 60.72 | correct |
| | 93 | 93.00 | 97.25 | 249 | 42.56 | correct |
| | 92 | 91.99 | 98.22 | 103 | 27.61 | correct |
| | 91 | 90.99 | 98.48 | 42 | 17.57 | correct |
| | 90 | 90.00 | 98.25 | 32 | 22.54 | correct |
| *Mycobacterium intracellulare* | 100 | 100.00 | 99.78 | 1241 | 66.33 | correct |
| | 99 | 98.92 | 98.92 | 1467 | 48.87 | correct |
| | 98 | 97.84 | 97.61 | 1796 | 59.83 | correct |
| | 97 | 96.98 | 98.26 | 460 | 15.32 | correct |
| | 96 | 95.91 | 97.84 | 984 | 32.77 | correct |
| | 95 | 94.85 | 99.14 | 959 | 31.93 | correct |
| | 94 | 93.98 | 98.28 | 1066 | 35.49 | correct |
| | 93 | 92.96 | 97.19 | 970 | 32.28 | correct |

| | | | | | | |
|---|---|---|---|---|---|---|
| | 92 | 91.86 | 98.06 | 683 | 22.64 | correct |
| | 91 | 90.99 | 96.54 | 1094 | 36.00 | correct |
| | 90 | 89.94 | 96.54 | 1249 | 40.99 | correct |
| *Pseudomonas aeruginosa* | 100 | 100.00 | 100.00 | 2822 | 99.44 | correct |
| | 99 | 98.94 | 99.79 | 2415 | 97.97 | correct |
| | 98 | 97.88 | 99.79 | 2125 | 93.16 | correct |
| | 97 | 96.84 | 98.72 | 1310 | 91.67 | correct |
| | 96 | 96.00 | 94.90 | 382 | 74.76 | correct |
| | 95 | 94.98 | 93.63 | 169 | 68.15 | correct |
| | 94 | 93.95 | 95.33 | 39 | 43.33 | correct |
| | 93 | 92.92 | 95.71 | 40 | 58.82 | correct |

### Table S3

**Table S3.** Third-party software utilised in the *LORCAN* pipeline.

| Program | Version | Author | Source |
|---|---|---|---|
| SeqKit | v. 0.8.0 | (Shen, Le et al. 2016) | https://github.com/shenwei356/seqkit |
| Porechop | v. 0.2.3 | Ryan Wick, University of Melbourne, Australia | https://github.com/rrwick/Porechop |
| minimap2 | v.2.5 | (Li 2018) | https://github.com/lh3/minimap2 |
| SAMtools | v.1.4 | (Li, Handsaker et al. 2009) | https://github.com/samtools/samtools |
| BLASTN | v.2.6.0 | (Altschul, Gish et al. 1990) | ftp://ftp.ncbi.nlm.nih.gov/blast/executables/blast+/LATEST/ncbi-blast-2.6.0+-x64-linux.tar.gz |
| MAFFT | v7.313 (2017/Nov/15) | (Katoh and Standley 2013) | http://mafft.cbrc.jp/alignment/software/ |
| Gblocks | GBLOCKS 0.91b | (Castresana 2000, Talavera and Castresana 2007) | http://molevol.cmima.csic.es/castresana/Gblocks.html |
| IQ-TREE | version 1.6.9 for Linux 64-bit | (Nguyen, Schmidt et al. 2014) | http://www.iqtree.org/ |
