## Supplementary material for "A sample-to-report solution for taxonomic identification of cultured bacteria in the clinical setting based on nanopore sequencing": SupportingTextS1.pdf

**Barcoding PCR Protocol:** based on **1D PCR barcoding (96) amplicons (SQK-LSK109),** Version: PBAC96_9069_v109_revK_14Aug2019 – Oxford Nanopore Technologies including all modifications made by the authors of this study (tested for amplicon lengths of 0.5 – 1 kb). Recommendations for long amplicons are based on the manufacturers' recommendations.

### Barcoding

- In a 0.2 ml 8-strip PCR tube, set up a barcoding PCR reaction as follows for each library:
    - 1 µl PCR Barcodes (one of BC1-BC96, at 10 µM)
    - 24 µl 1.05 nM first-round PCR product
    - 25 µl Taq 2x master mix (LongAmp Taq 2x master mix for longer amplicons)
- Mix by pipetting.
- Seal the tubes and briefly spin down.
- Amplify using the following cycling conditions (adjust numbers of cycles and extension time for amplicons > 1kb):
    - Initial denaturation 3 mins @ 95 °C, 1 cycle
    - Denaturation 15 secs @ 95 °C, 12 cycles
    - Annealing 15 secs @ 62 °C, 12 cycles
    - Extension 1 min @ 65 °C, 12 cycles
    - Final extension @ 65 °C, 1 cycle
    - Hold @ 4 °C
- Quantify each raw barcoding PCR product with a fluorometric method (tested with Qubit BR assay).
- Combine the barcoded libraries in the desired ratios in a 1.5 ml reaction tube.
- SPRI-bead clean-up (tested with AMPure XP and CleanNGS beads):
    - Bring the SPRI-beads to room temperature; resuspend by vortexing.
    - Add 0.8 volumes resuspended SPRI-beads to 1 volume of library mix and mix by flicking the tube (adjust ratios for longer amplicons in necessary).
    - Incubate on a Hula mixer (rotator mixer) for 5 minutes at RT.
    - Prepare 1 ml of fresh 70% ethanol in Nuclease-free water.
    - Spin down the sample and pellet on a magnet. Keep the tube on the magnet, and pipette off the supernatant.
    - Keep the tube on the magnet, wash beads with 400 µl of freshly prepared 70% ethanol without disturbing the pellet.
    - Remove and discard the 70% ethanol using a pipette.
    - Repeat the previous washing step.
    - Spin down and place the tube back on the magnet. Pipette off any residual ethanol. Allow to dry for ~30 seconds, but do not dry the pellet to the point of cracking.

- Remove the tube from the magnetic rack and resuspend the pellet in 50 µl Nuclease-free water. Incubate for 2 minutes at RT.
    - Pellet the beads on a magnet until the eluate is clear and colourless.
    - Remove and retain 48 µl of eluate into a clean 1.5 ml reaction tube.
- Quantify the barcoded library using a fluorometric method.
- Dilute the library pool if the total concentration exceeds 140 nM (tested range: 60-140 nM).
- Proceed to End-Repair.

Example: pooling of barcoding-PCR products

| Barcode | DNA-Nr. | Assay (Primer set) | Fragment-length (bp) | Concentration [ng/µl] | Concentration [pmol/µl] | DNA required [pmol] | Volume to create the pooled library [µl] |
|---|---|---|---|---|---|---|---|
| 01 | sample 1 | Mycobacteria | 1000 | 17 | 0.0262 | 1.0256 | 39.2 |
| 02 | sample 2 | general bacteria | 500 | 15 | 0.0462 | 1.0256 | 22.2 |
| 03 | sample 3 | general bacteria | 500 | 21 | 0.0646 | 1.0256 | 15.9 |
| 04 | sample 4 | general bacteria | 500 | 22 | 0.0677 | 1.0256 | 15.2 |
| 05 | sample 5 | Mycobacteria | 1000 | 15 | 0.0231 | 1.0256 | 44.4 |
| 06 | positive control | Mycobacteria | 500 | 20 | 0.0615 | 1.0256 | 16.7 |
| 91 | place-holder | general bacteria | 500 | 20 | 0.0615 | 1.0256 | 16.7 |
| 91 | place-holder | general bacteria | 500 | 20 | 0.0615 | 1.0256 | 16.7 |
| 91 | place-holder | general bacteria | 500 | 20 | 0.0615 | 1.0256 | 16.7 |

**DNA repair and end-prep**

Prepare the reagents by placing both NEBNext FFPE DNA Repair Mix and NEBNext End repair / dA-tailing Module on ice, bring both Ultra II End-prep reaction buffer and NEBNext FFPE DNA Repair buffer to room temperature, and verify that the tubes do not contain precipitates. If precipitates are visible incubate the buffer at 37°C and mix well until precipitate disappears.

- In a 1.5 ml tube mix the following:
  - 1 µl NFW
  - 3.5 µl NEBNext FFPE DNA Repair buffer
  - 2 µl NEBNext FFPE DNA Repair Mix
  - 3.5 µl Ultra II End-prep reaction buffer
  - 3 µl Ultra II End-prep enzyme Mix
  - 47 µl of purified library
- Mix gently by flicking the tube, transfer to a 0.2 ml PCR tube and spin down.
- Using a thermal cycler, incubate at 20° C for 5 min, and 65° C for 5 min.
- SPRI-bead clean-up (tested with AMPure XP and CleanNGS beads):
  - Bring the SPRI-beads to room temperature; resuspend by vortexing.
  - Transfer the DNA sample to a clean 1.5 ml reaction tube.
  - Add 60 µl of resuspended SPRI-beads to the end-prep reaction and mix by flicking the tube.
  - Incubate on a Hula mixer (rotator mixer) for 5 minutes at RT.
  - Prepare 500 µl of fresh 70% ethanol in Nuclease-free water.
  - Spin down the sample and pellet on a magnet. Keep the tube on the magnet, and pipette off the supernatant.
  - Keep the tube on the magnet, wash beads with 200 µl of freshly prepared 70% ethanol without disturbing the pellet.
  - Remove the 70% ethanol using a pipette and discard.
  - Repeat the previous washing step.
  - Spin down and place the tube back on the magnet. Pipette off any residual ethanol. Allow to dry for ~30 seconds, but do not dry the pellet to the point of cracking.
  - Remove the tube from the magnetic rack and resuspend the pellet in 61 µl of Nuclease-free water. Incubate for 2 minutes at RT.
  - Pellet the beads on a magnet until the eluate is clear and colourless.
  - Remove and retain 61 µl of eluate into a clean 1.5 ml reaction tube.
  - Quantify 1 µl of eluted sample using a fluorometric method.
  - Proceeded to the adapter ligation step or store the sample at 4° C overnight.

**Adapter ligation and clean-up**

- Spin down Adapter Mix (AMX) and T4 Ligase, and place on ice.
- Thaw Ligation Buffer (LNB) at RT, spin down and mix by pipetting. Due to viscosity, vortexing this buffer is ineffective. Place on ice immediately after thawing and mixing.
- Thaw the Elution Buffer (EB) at RT, mix by vortexing, spin down.
- To retain DNA fragments of all sizes, thaw one tube of Short Fragment Buffer (SFB) at RT, mix by vortexing, spin down (substitute with of Long Fragment Buffer (LFB) for fragment sizes of >3 kb).
- In a 1.5 ml reaction tube, mix in the following order:
  - 60 µl library-mix from the previous step
  - 25 µl Ligation Buffer (LNB)
  - 10 µl NEBNext Quick T4 DNA Ligase
  - 5 µl Adapter Mix (AMX)
- Mix gently by flicking the tube, and spin down.
- Incubate the reaction for 10 minutes at RT.
- SPRI-bead clean-up (tested with AMPure XP and CleanNGS beads):
  - Bring the SPRI-beads to room temperature; resuspend by vortexing.
  - Add 40 µl of resuspended SPRI-beads to the reaction and mix by flicking the tube.
  - Incubate on a Hula mixer (rotator mixer) for 5 minutes at RT.
  - Spin down the sample briefly and pellet on a magnet. Keep the tube on the magnet, and pipette off the supernatant.
  - Add 250 µl Short Fragment Buffer (SFB) for washing (substitute with Long Fragment Buffer (LFB) for long fragments).
  - Flick the beads to resuspend, then return the tube to the magnetic rack and allow the beads to pellet.
  - Remove and discard the supernatant using a pipette.
  - Repeat the previous step.
  - Spin down briefly and place the tube back on the magnet. Pipette off any residual supernatant. Allow to dry for approx. 30 seconds, but do not dry the pellet to the point of cracking.
  - Remove the tube from the magnetic rack and resuspend pellet in 15 µl Elution Buffer (EB). Incubate for 10 minutes at RT. (For high molecular weight DNA, incubation at 37° C can improve the recovery of long fragments).
  - Pellet the beads on a magnet until the eluate is clear and colourless.
  - Transfer 15 µl of eluate into a clean 1.5 ml reaction tube.
  - Dispose of the pelleted beads.
  - Quantify 1 µl of eluted sample using a fluorometric method.
- Store the library on ice until proceeding to the next steps.

### Priming and loading the SpotON flow cell

Priming and loading of the flowcells was performed according to the manufacturer's instructions.

### Reagents and materials

| Name | Product code | Manufacturer |
|---|---|---|
| Barcode Mix BC01 bis BC96 (weisse Kappen) | EXP-PBC096 | Oxford Nanopore Tech. (OX, UK) |
| Adapter Mix | SQK-LSK109 | Oxford Nanopore Tech. |
| Ligation Buffer | SQK-LSK109 | Oxford Nanopore Tech. |
| Elution buffer | SQK-LSK109 | Oxford Nanopore Tech. |
| Small Fragment Buffer | SQK-LSK109 | Oxford Nanopore Tech. |
| Sequencing Buffer | SQK-LSK109 | Oxford Nanopore Tech. |
| Loading Beads | SQK-LSK109 | Oxford Nanopore Tech. |
| Flush Tether | SQK-LSK109 | Oxford Nanopore Tech. |
| Flush Buffer | SQK-LSK109 | Oxford Nanopore Tech. |
| Flowcell SpotON Mk I (R9.4) | FLO-MIN106.1 | Oxford Nanopore Tech. |
| 1.5 ml DNA LoBind Tubes | 30108051 | Eppendorf (Hamburg, DE) |
| 8-Tube PCR strips | XT90.1 | Brand (Wertheim, DE) |
| NEBNext End Repair / dA-tailing Module | E7546S | New England Biolabs (ON, CA) |
| NEBNext FFPE DNA Repair Mix | M6630S | New England Biolabs |
| NEBNext Quick Ligation Module | E6056S | New England Biolabs |
| Taq 2X Master Mix | M0270 | New England Biolabs |
| CleanNGS DNA Beads | CNGS0005 | CleanNA (Waddinxveen, NL) |
| Ethanol BioUltra ≥99.8% | 51976 | Sigma-Aldrich (Merck, Darmstadt, DE) |
| Nuclease free water (NFW) | AM9937 | Ambion (Thermo Fisher Scientific Inc., MA, USA) |
| Solution A | EXP-WSH002 | Oxford Nanopore Tech. |
| Storage Buffer | EXP-WSH002 | Oxford Nanopore Tech. |
| Qubit dsDNA BR Assay | Q32850 | Invitrogen (CA, USA) |
| PCR-Tubes, flat caps, clear 0,5 ml | 732-0675 | Axygen (Corning, NY USA) |
